# The half-life of the bone-derived hormone osteocalcin is regulated through *O*-glycosylation in mice, but not in humans

**DOI:** 10.1101/2020.07.16.206656

**Authors:** Omar Al Rifai, Catherine Julien, Denis Faubert, Erandi Lira-Navarrete, Yoshiki Narimatsu, Julie Lacombe, Henrik Clausen, Mathieu Ferron

## Abstract

Osteocalcin (OCN) is an osteoblast-derived hormone with pleiotropic physiological functions. Like many peptide hormones, OCN is subjected to post-translational modifications (PTMs) which control its activity. Here, we uncover *O*-glycosylation as a novel PTM present on mouse OCN and occurring on a single serine (S8) independently of its carboxylation and endoproteolysis, two other PTMs regulating this hormone. We also show that *O*-glycosylation increases OCN half-life in plasma ex vivo and in the circulation in vivo. Remarkably, in human OCN (hOCN), the residue corresponding to S8 is a tyrosine (Y12), which is not *O-*glycosylated. Yet, the Y12S mutation is sufficient to *O*-glycosylate hOCN and to increase its half-life in plasma compared to wildtype hOCN. These findings reveal an important species difference in OCN regulation, which may explain why serum concentrations of OCN are higher in mouse than in human.

The authors have nothing to disclose

## INTRODUCTION

Osteocalcin (OCN) is a peptide hormone secreted by osteoblasts, the bone forming cells. It regulates glucose metabolism by promoting beta cells proliferation and insulin secretion, and by improving insulin sensitivity (Pi et al., 2011, Ferron et al., 2012). In addition to its role in the regulation of energy metabolism, OCN is also involved in male fertility by promoting testosterone synthesis by Leydig cells (Oury et al., 2011), in muscle adaptation to exercise by improving glucose and fatty acid uptake in myocytes (Mera et al., 2016a), and in acute stress response through the inhibition of post-synaptic parasympathetic neurons (Berger et al., 2019). Overall, OCN might acts as an “anti-geronic” circulating factor preventing age-related cognitive decline and muscle wasting (Khrimian et al., 2017, Mera et al., 2016b, Oury et al., 2013b). The G protein coupled receptor family C group 6 member A (GPRC6A) mediates OCN function in beta cells, muscles and testis (Mera et al., 2016a, Pi et al., 2011, Oury et al., 2013a), while the G protein coupled receptor 158 (Gpr158) mediates its function in the brain (Khrimian et al., 2017, Kosmidis et al., 2018).

Within the bone tissue, OCN undergoes a series of post-translational modifications (PTM) that are critical for the regulation of its endocrine functions. Prior to its secretion, in the osteoblast endoplasmic reticulum, the OCN precursor (pro-OCN) is γ-carboxylated on three glutamic acid residues (Glu) by the vitamin K-dependent γ-glutamyl carboxylase (Ferron et al., 2015). In the trans-Golgi network, pro-OCN is next cleaved by the proprotein convertase furin releasing mature carboxylated OCN (Gla-OCN) (Al Rifai et al., 2017). The presence of the negatively charged Gla residues allows Gla-OCN to bind hydroxyapatite, the mineral component of the bone extracellular matrix (ECM). It is during bone resorption that Gla-OCN is decarboxylated through a non-enzymatic process involving the acidic pH generated by the osteoclasts, ultimately leading to the release of bioactive uncarboxylated OCN (ucOCN) in the circulation (Ferron et al., 2010a, Lacombe et al., 2013). The conclusion that ucOCN represents the bioactive form of this protein in rodents is supported by cell-based assays, mouse genetics and in vivo studies [reviewed in (Mera et al., 2018)].

The role of OCN in the regulation of glucose metabolism appears to be conserved in humans. Human ucOCN can bind and activate human GPRC6A (De Toni et al., 2016) and promotes beta cells proliferation and insulin synthesis in human islets (Sabek et al., 2015), while mutations or polymorphisms in human *GPRC6A* are associated with insulin resistance (Di Nisio et al., 2017, Oury et al., 2013a). Finally, several cross-sectional and observational studies have detected a negative association between OCN or ucOCN, and insulin resistance or the risk of developing type 2 diabetes in various human populations (Lin et al., 2018, Turcotte et al., 2020, Lacombe et al., 2020).

Yet, some important species divergences exist between mice and humans with regard to OCN biology. First, only 30 out of the 46 amino acids (i.e., 65%) composing mature mouse OCN are conserved in human OCN. This is in striking contrast with other peptide hormones involved in the control of energy metabolism such as leptin and insulin whose respective sequence display about 85% conservation between mouse and human. Second, the circulating concentrations of OCN, even though decreasing with age in both species, are five to ten times higher in mice than in humans throughout life span [(Mera et al., 2016a) see also Supplemental table 1]. Based on these observations, we hypothesized that the posttranslational regulation of OCN may be different between these two species, resulting in increased half-life of mouse OCN in circulation.

Here, using proteomics and cell-based assays, we identified *O*-glycosylation as a novel PTM presents in mouse OCN, and showed that this modification increases mouse OCN half-life in plasma ex vivo and in vivo. In contrast, mature human OCN does not contain the *O*-glycosylation site found in the mouse protein and consequently is not normally glycosylated. Yet, a single point mutation in human OCN is sufficient to elicit its *O*-glycosylation and to increase its half-life in plasma.

## RESULTS

### Mouse OCN is *O-*glycosylated on a single serine residue

To better document the circulating level of OCN in humans and mice, we measured the serum concentration of OCN in wildtype mice at different age (2 to 60 weeks) and compared the values with the reported serum level of OCN at corresponding life periods in humans (Supplemental Table 1). This analysis reveals that serum OCN level is five- to ten-time lower in humans than in mice throughout life. One potential explanation for this observation could be that a mouse specific PTM increases mouse OCN half-life in circulation. Since OCN Gla residues and pro-OCN cleavage site are conserved between mouse and human, we searched for additional PTMs present in mouse OCN and characterized their impact on OCN half-life.

To that end, OCN was immunoprecipitated from the secretion media of primary mouse osteoblast cultures or from mouse bone protein extracts using specific polyclonal goat antibodies recognizing OCN C-terminus sequence (Ferron et al., 2010b). OCN was then characterized without proteolysis by reverse-phase HPLC followed by mass spectrometry (MS) and tandem mass spectrometry (MS/MS). This “top-down” analysis revealed that the most abundant OCN forms have a monoisotopic mass ranging from 5767.6961 to 6441.7636 Da which exceeds the predicted mass of 5243.45 Da corresponding to the fully carboxylated (3×Gla) OCN (Supplemental Table 2 and 3). According to the various monoisotopic masses observed, we predict that this difference could be mainly explained by the presence of a single *O-*linked glycan adduct composed of one *N*-acetylgalactosamine (GalNAc), one galactose (Gal) and one or two *N*-acetylneuraminic acid (NANA). In addition to *O*-glycosylation, minor additional monoisotopic mass change corresponding to oxidation, unidentified modifications and/or adduct ions were also detected. Uncarboxylated OCN does not accumulate in bone ECM (Ferron et al., 2015) and accordingly, only fully or partially carboxylated OCN was detected in bone extracts, both of which were found to be *O*-glycosylated. In contrast, significant amount of ucOCN could be detected in osteoblast culture supernatants, most likely due to the low level of vitamin K present fetal bovine serum. Importantly, in osteoblast supernatant the *O-*linked glycan adduct could be detected on both carboxylated and uncarboxylated OCN (Supplemental Table 2).

Using multiple approaches, we next established that mouse OCN is indeed subjected to *O-*glycosylation in cells and in vivo. First, in SDS-PAGE analyses the apparent molecular weight of OCN is reduced when expressed in HEK293 cells where the O-glycosylation capacity has been engineered to truncate O-glycans by knockout of *COSMC* (core 1β3-Gal-T-specific molecular chaperone), a private chaperone required for the elongation of *O*-glycans (Figure 1A) (Steentoft et al., 2011). Second, when expressed in CHO-ldlD cells, which have defective UDP-Gal/UDP-GalNAc 4-epimerase and are hence deficient in *O*-glycosylation (Kingsley et al., 1986), OCN apparent molecular weight is also reduced compared to the same protein expressed in the parental CHO cell line. Supplementation of Gal and GalNAc in the culture media rescued the *O-*glycosylation defect of CHO-ldlD and restored the molecular shift in the secreted OCN (Figure 1B). Third, treatment of primary osteoblasts with benzyl-N-acetyl-α-galactosaminide (GalNAc-bn), an inhibitor of *N*-acetylgalactosaminyltransferases (GalNAc-Ts), the enzymes responsible for initiating *O-*glycosylation, decreases the apparent molecular weight of OCN secreted in the media (Figure 1C). Finally, treatment of mouse bone extracts with neuraminidase and *O*-glycosidase, which removes respectively NANA, and core 1 and core 3 *O*-linked disaccharides, also decreases the apparent molecular weight of endogenous OCN (Figure 1D).

**Figure 1.**
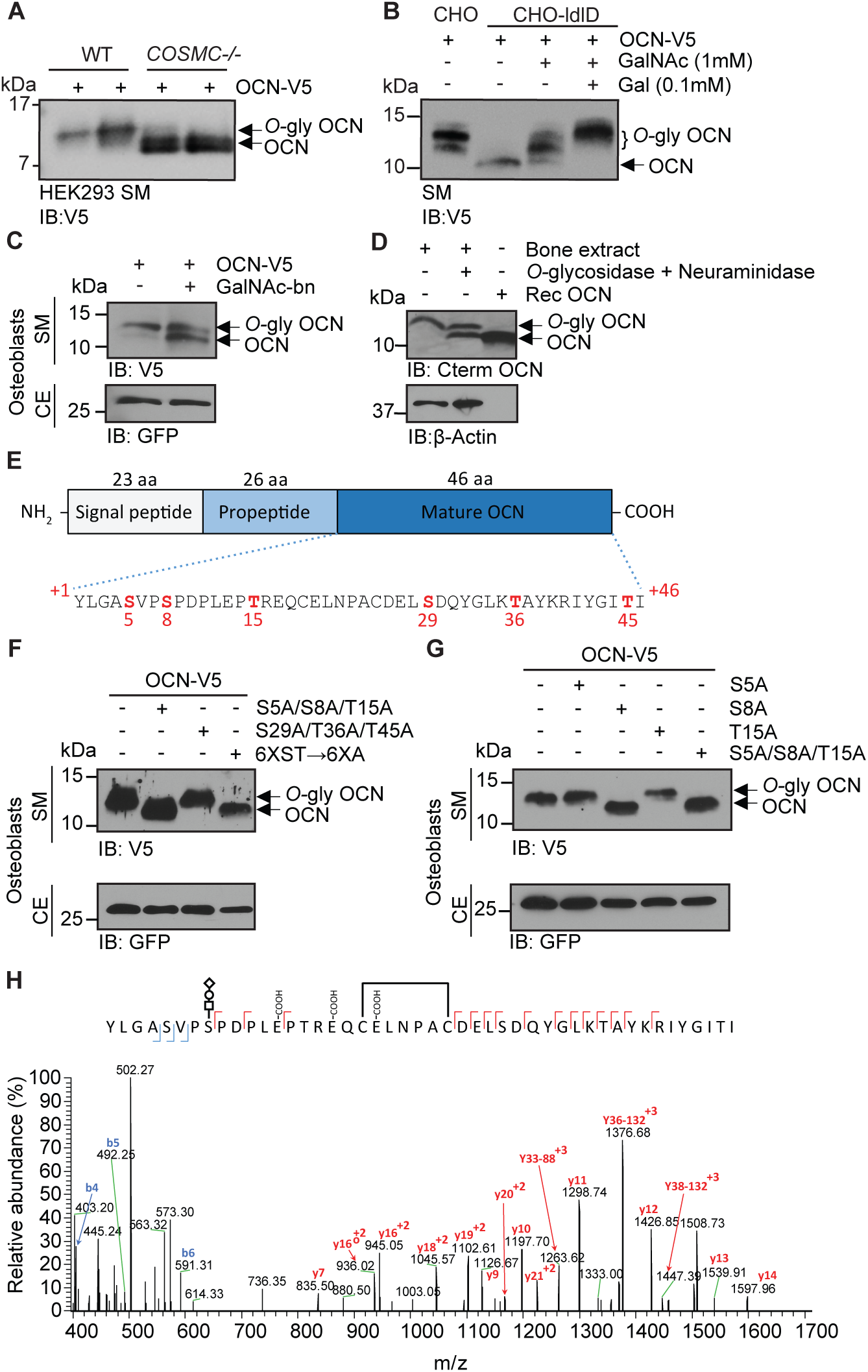
OCN is O-glycosylated in vitro and in vivo on serine 8. **(A)** Western blot analysis on the secretion media (SM) of HEK293 (WT) and HEK293 lacking COSMC (COSMC-/-) transfected with mouse OCN-V5. **(B)** Western blot analysis on the secretion media (SM) of CHO and CHO-ldlD cells transfected with mouse OCN-V5. CHO-ldlD cells were treated or not with 0.1 mM Galactose (Gal) and/or 1 mM N-acetylgalactosamine (GalNAc). **(C)** Effect of N-acetylgalactosaminyltransferase (GalNAc-Ts) inhibition on mouse OCN O-glycosylation in osteoblasts. Western blot analysis on the secretion media (SM) and cell extract (CE) of primary osteoblasts transfected with mouse OCN-V5 and treated or not with 2 mM of GalNAc-bn. **(D)** OCN deglycosylation assay. Bone extract of C57BL/6J mice were treated or not with O-glycosidase and neuraminidase for 4 hours at 37°C and analyzed by Western blot using anti-C-termimal OCN antibody (Cterm OCN). β-actin was used as a loading control. Rec OCN: Non-glycosylated OCN produced in bacteria. **(E)** Structure of mouse pre-pro-OCN and amino acid sequence of mature mouse OCN. The serine (S) and threonine (S) residues are in red. **(F)** Western blot analysis on the secretion media (SM) and cell extract (CE) of primary osteoblasts transfected with OCN-V5 containing or not the indicated mutations. In the 6XST→6XA mutant, all six serine and threonine residues from OCN were mutated to alanine. **(G)** Western blot analysis on the secretion media (SM) and cell extract (CE) of primary osteoblasts transfected with OCN-V5 containing or not the indicated mutations. **(H)** Annotated HCD MS/MS spectrum of a modified form of OCN (HexNAc-Hex-NANA + 3 Gla + S-S) pulled down from the bone homogenate of C57BL/6J mice. The precursor m/z value is 1180.95003 (M+5H)^+5^ and mass accuracy with the annotated OCN modified form is 4.6 ppm. In **C, F** and **G**, GFP co-expressed from OCN-V5 expression vector, was used as a loading control.

We next aimed at identifying which OCN residue(s) is(are) *O*-glycosylated. Mature mouse OCN contains 3 serine (S) and 3 threonine (T) residues (Figure 1E), the two main types of amino acids on which *O-*glycosylation occurs (Steentoft et al., 2013). As expected, mutating all serine and threonine residues into alanine abrogates OCN glycosylation in primary mouse osteoblasts as assessed by SDS-PAGE (Figure 1F). Further mutagenesis studies revealed that the *O-*glycosylation site resides within the N-terminal part of the protein, i.e., on S5, S8 or T15 (Figure 1F). Single amino acid mutagenesis allowed the identification of S8 as the *O-*glycosylation site of OCN in osteoblasts (Figure 1G), a result consistent with the MS/MS analysis of OCN isolated from bone which also suggested that this residue is the *O*-glycosylation site (Figure 1H). Together, these results establish that mouse OCN is *O-*glycosylated on at least one serine residue in cell culture and in vivo.

### Several polypeptide *N*-acetylgalactosaminyltransferases (GalNAc-Ts) redundantly *O*-glycosylate OCN independently of its carboxylation and processing

Protein *O*-glycosylation is initiated by the transfer of a GalNAc to a serine or threonine residue, a reaction taking place in the Golgi and catalyzed by GalNAc-Ts, a family of enzymes comprising 19 different members in mice (Bennett et al., 2012). Quantitative PCR on mRNA isolated from undifferentiated and differentiated primary mouse osteoblasts revealed that several GalNAc-Ts are expressed in this cell type, with *Galnt1* and *Galnt2* being the most strongly expressed ones (Figure 2A). We noticed that S8A mutation abrogates OCN *O*-glycosylation in HEK293 cells and in primary osteoblasts (Figure 1G and data not shown). GalNAc-T3 and its paralogue GalNAc-T6 are known to be expressed in HEK293 (Narimatsu et al., 2019b) and our data shows they are also expressed in primary mouse osteoblasts and induced during osteoblast differentiation. Although these observations suggest one or both of these enzymes may be involved in OCN *O*-glycosylation, the inactivation of *GALNT3* and/or *GALNT6* genes failed to alter OCN *O*-glycosylation in HEK293 (Figure 2B). Since *GALNT1* and *GALNT2* are also highly expressed in osteoblasts and induced during osteoblast differentiation, we inactivated these two genes in combination with *GALNT3*, and assess the impact on OCN *O*-glycosylation in HEK293 cells. This partially abolished OCN glycosylation (Figure 2B), suggesting that these three GalNAc-Ts are the primary isoenzymes that redundantly initiate the *O*-glycosylation of OCN.

**Figure 2.**
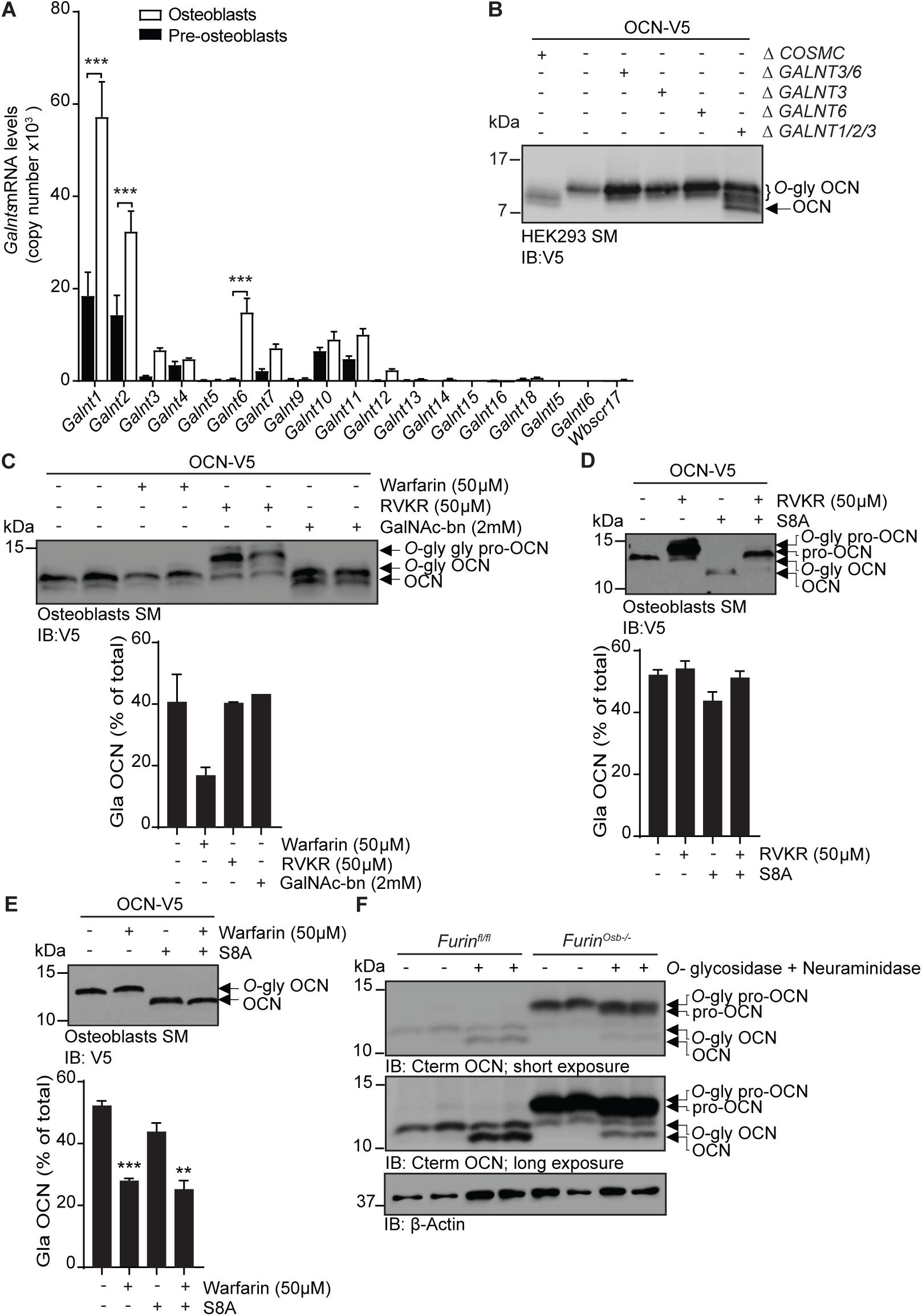
OCN O-glycosylation by N-acetylgalactosaminyltransferase (GalNAc-Ts) is independent of its processing and γ-carboxylation. **(A)** Galnts expression in pre-osteoblasts (undifferentiated) and osteoblasts (differentiated) by quantitative PCR. Results are represented as copy number of Galnts normalized to Actb. **(B)** Western blot analysis of OCN in the secretion media (SM) of HEK293 cells deficient for specific GalNAc-Ts. OCN-V5 was transfected in parental, COSMC-/- (ΔCOSMC), or GALNTs deficient (Δ) HEK293 cells and analysed by Western blot using anti-V5 antibody. **(C)** Western blot analysis on the secretion media (SM) of osteoblasts transfected with mouse OCN-V5 and treated or not with 2 mM of GalNAc-bn, 50 μM warfarin or 50 μM Dec-RVKR-CMK (RVKR) (upper panel), and percentage of carboxylated OCN (Gla OCN) over total OCN measured by ELISA (lower panel). **(D)** Western blot analysis on the secretion media (SM) of osteoblasts transfected with mouse OCN-V5 containing or not the S8A mutation and treated with 50 μM Dec-RVKR-CMK (RVKR) (upper panel), and percentage of carboxylated OCN (Gla OCN) over total OCN measured by ELISA (lower panel). **(E)** Western blot analysis on the secretion media (SM) of osteoblasts transfected with mouse OCN-V5 containing or not the S8A mutation and treated with 50 μM warfarin (upper panel), and percentage of carboxylated OCN over total OCN measured by ELISA (lower panel). **(F)** Western blot analysis of OCN deglycosylation assay on bone extracts from Furin^fl/fl^ and Furin^Osb-/-^ mice. Bone extracts were treated or not with O-glycosidase and neuraminidase for 4 hours at 37°C and analyzed by Western blot using anti-C-termimal OCN antibody (Cterm OCN). ** p<0.01; *** p<0.001 using one-way ANOVA with Bonferroni multiple comparisons test.

Processing of pro-OCN by the proprotein convertase furin and its γ-carboxylation are two post-translational modifications regulating OCN endocrine function (Ferron et al., 2015, Al Rifai et al., 2017). We therefore next aimed at testing whether OCN *O-*glycosylation can interfere with its γ-carboxylation or processing, or inversely, if *O*-glycosylation is modulated by γ-carboxylation or processing of pro-OCN. Pharmacological inhibition of γ-carboxylation or furin, using warfarin or Dec-RVKR-CMK (RVKR) respectively, did not impact OCN *O-*glycosylation in primary osteoblasts and HEK293 cells (Figure 2C and Supplemental figure 1A). Similarly, inhibition of OCN *O-*glycosylation through GalNAc-bn treatment or the S8A mutation did not significantly affect its processing or its γ-carboxylation (Figure 2C-E and Supplemental figure 1A). We also tested whether OCN processing influences its *O-*glycosylation in vivo. As shown in Figure 2F, both mature OCN present in control bones and pro-OCN present in furin-deficient bones are de-glycosylated by neuraminidase and *O*-glycosidase, indicating that pro-OCN is normally *O-*glycosylated in absence of processing by furin in vivo. Altogether, these results support the notion that osteocalcin *O-*glycosylation is not affected by its carboxylation status or by its processing by furin. Moreover, blocking *O-*glycosylation does not prevent pro-OCN processing by furin or its carboxylation.

### *O*-glycosylation increases mouse OCN half-life in plasma ex vivo by preventing plasmin-mediated endoproteolysis

The results presented above suggest that *O-*glycosylation is not regulating the processing of pro-OCN by furin or the secretion of mature OCN by osteoblasts. It was recently observed, that *O*-glycosylation can also increase the stability of some peptide hormones in the circulation by preventing proteolytic degradation (Hansen et al., 2019, Madsen et al., 2020). We therefore aimed at testing the impact of *O*-glycosylation on OCN half-life in plasma. To that end, we produced and purified *O*-glycosylated ucOCN from HEK293 and first compared its purity and molecular weight to native non-glycosylated ucOCN produced in bacteria by LC-MS, LC-MS/MS and SDS-PAGE (Supplemental Figure 2A, B; Figure 3A, B). Importantly, we observed that >99% of the ucOCN purified from HEK293 is O-glycosylated, with a certain proportion (∼30%) containing two glycan adducts (Figure 3A), suggesting that in this context O-glycosylation may occurs on more than one residues. Freshly isolated *Ocn*-/- plasma, which is depleted of endogenous OCN, was next used to assess OCN half-life ex vivo. Non-glycosylated ucOCN has a half-life of ∼120 minutes when incubated in plasma at 37°C, while *O*-glycosylated ucOCN is stable for more than 5 hours in the same conditions (Figure 3C). Non-glycosylated ucOCN was stable when incubated in a saline solution containing 3.5% BSA at 37°C for 2 hours (Supplemental Figure 2C), implying that ucOCN is not intrinsically unstable. In addition, stability of the non-glycosylated ucOCN was restored when incubated in plasma at 4°C or in heat-inactivated (HI) plasma at 37°C (Figure 3D), suggesting that non-glycosylated ucOCN’s decline involves the action of a protease. Pepstatin A (Pep A) an aspartic

**Figure 3.**
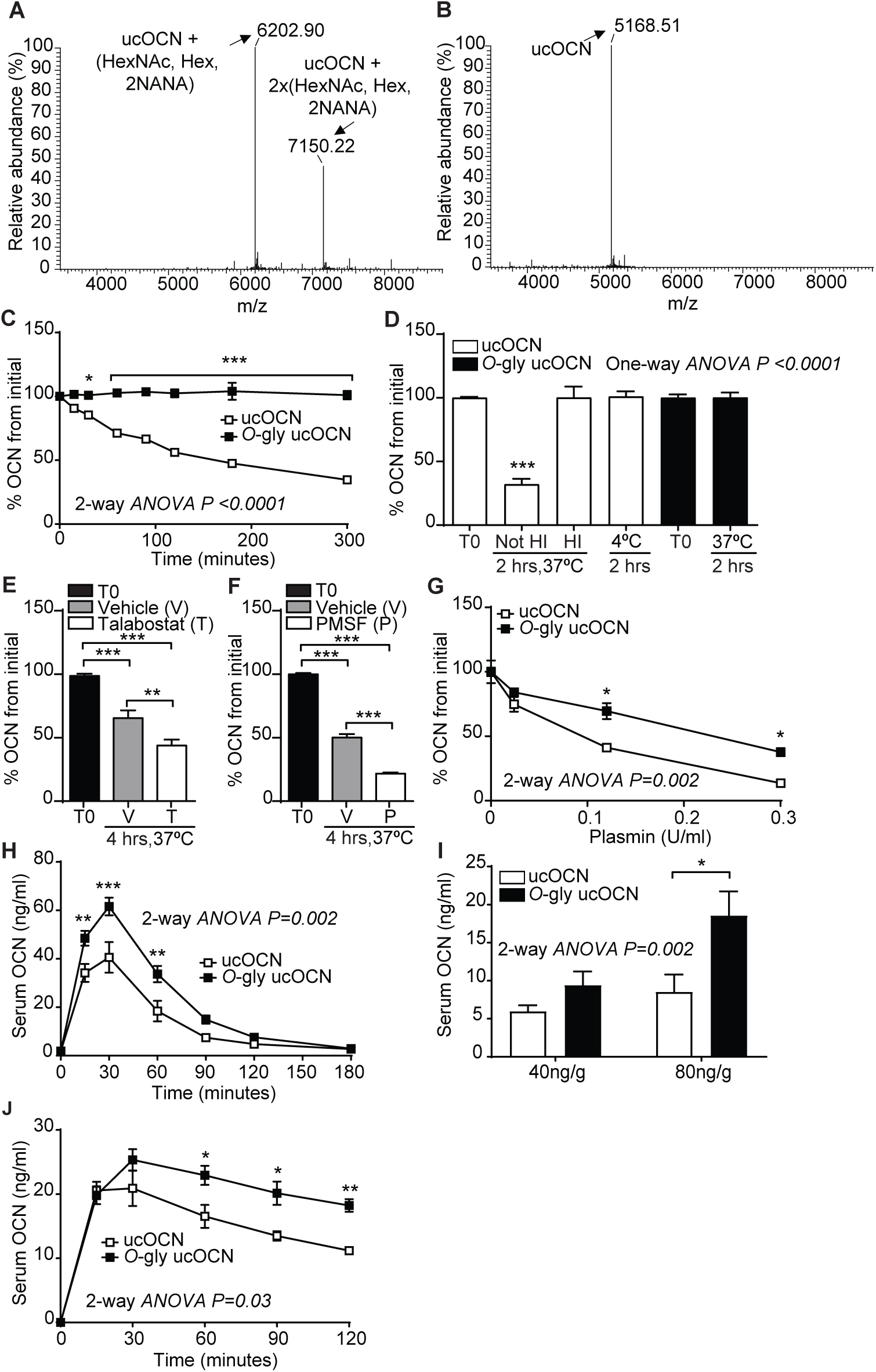
Mouse OCN O-glycosylation increases its half-life ex vivo and in vivo. **(A)** Annotated and deconvoluted MS spectrum of purified glycosylated mouse OCN (O-gly ucOCN). **(B)** Annotated and deconvoluted MS spectrum of purified non-glycosylated mouse OCN (ucOCN). **(C-D)** Ex vivo half-life of O-gly ucOCN and ucOCN in OCN deficient (Ocn-/-) plasma (n=3-4 plasma). **(C)** 100 ng/ml of O-gly ucOCN or ucOCN were incubated in plasma at 37°C for 0 to 5 hours and OCN levels were measured at the indicated times. **(D)** 100 ng/ml of O-gly ucOCN or ucOCN were incubated in plasma for 2 hours at 37°C in different conditions or at 4°C. HI: heat inactivated plasma. **(E)** 50 ng/ml of ucOCN was incubated in plasma from Ocn-/- mice for 4 hours at 37°C in the presence of vehicle (V) or 10 mM talabostat (T). **(F)** 50 ng/ml of ucOCN was incubated in plasma from Ocn-/- mice for 4 hours at 37°C in the presence of vehicle (V) or 10 mM phenylmethylsulfonyl fluoride (PMSF). **(G)** Effect of plasmin on OCN stability ex vivo. 50 ng/ml of O-gly ucOCN and ucOCN were incubated for two hours in Ocn-/- heat-inactivated plasma containing different concentration of plasmin. **(H)** In vivo half-life of O-gly ucOCN and ucOCN in fed condition in mice. O-gly ucOCN or ucOCN were injected intraperitoneally in OCN deficient male mice (Ocn-/-; n=9 mice each) at a dose of 40ng/g of body weight and serum OCN levels were measured at the indicated time points. **(I)** O-gly ucOCN (n=4 mice) or ucOCN (n=4 mice) were injected intraperitoneally in OCN deficient male mice (Ocn- /-), in fed condition, at a dose of 40 or 80ng/g of body weight. Serum OCN level was measured two hours post-injection. **(J)** In vivo half-life of O-gly ucOCN and ucOCN in fasting condition. OCN deficient male mice (Ocn-/-) were fasted for 16 hours, O-gly ucOCN (n=5 mice) or ucOCN (n=5 mice) were injected intraperitoneally at a dose of 40ng/g of body weight and serum OCN levels were measured at the indicated time points. T0: start point, see material and methods; HexNAc: N-acetylhexosamine; Hex: Hexose; NANA: N-acetylneuraminic acid. OCN measurements were performed using total mouse OCN ELISA assay (see Material and Methods). Results are given as mean ±SEM. * p<0.05; ** p<0.01; *** p<0.001 using 2-way ANOVA for repeated measurements with Bonferroni multiple comparisons test.

proteases inhibitor, RVKR a proprotein convertases inhibitor and ethylenediaminetetraacetic acid (EDTA) which inhibits metalloproteases did not affect non-glycosylated ucOCN stability in plasma ex vivo (Supplemental Figure 2C-D). The OCN sequence surrounding S8 contains several proline residues (Figure 1E) and could therefore be recognized by prolyl endopeptidases, such as the fibroblast activation protein (FAP) which is present in the circulation (Sanchez-Garrido et al., 2016, Coutts et al., 1996). We thus also tested the effect of talabostat, an inhibitor of FAP and dipeptidyl peptidases, on ucOCN stability in plasma. Surprisingly, treatment of plasma with 10 mM talabostat did not inhibit, but rather increased non-glycosylated ucOCN degradation (Figure 3E). A reduced stability of non-glycosylated OCN in plasma was also observed in the presence of PMSF (Figure 3F) at a concentration (10 mM) that was also shown to inhibit dipeptidyl peptidases and prolyl endopeptidases (Banbula et al., 2000, Bermpohl et al., 1998). One function of FAP in vivo is to activate the α2-antiplasmin precursor releasing the active α2-antiplasmin, which in turn acts as an inhibitor of plasmin activity (Lee et al., 2011, Lee et al., 2006). Interestingly, mouse OCN contains arginine residues in its N- and C-terminus (R16 and R40 respectively), which could be potential cleavage sites for plasmin (Rawlings et al., 2008). Together these observations led us to hypothesize that plasmin could be a protease responsible for ucOCN degradation in plasma. Supporting this notion, ucOCN is rapidly degraded when low concentration of recombinant plasmin is added to previously heat-inactivated plasma or in Tris buffered solution (Figure 3G and data not shown). Moreover, *O-*glycosylation partially protect ucOCN from plasmin-mediated degradation in the same assay (Figure 3G). Altogether, these data indicate that *O-*glycosylation protects ucOCN from plasmin mediated proteolysis, thereby increasing its half-life in plasma in vitro.

### Glycosylation increases mouse OCN stability in vivo

We next examined the stability of glycosylated and non-glycosylated mouse ucOCN in vivo by injecting an equal dose (40ng/g of body weight) of each of these proteins in *Ocn*- /- mice which are depleted of endogenous OCN. In fed mice, the maximum serum level of ucOCN reached 30 minutes after injection was 1.5 times higher with the glycosylated protein compared to the non-glycosylated form (Figure 3H). Moreover, glycosylated ucOCN serum concentration remains higher than the one of non-glycosylated ucOCN for up to 2 hours following the injection. Circulating level of glycosylated ucOCN after 2 hours was further increased when 80ng/g of body weight of protein was injected, while non-glycosylated ucOCN was not significantly increased with this higher dose (Figure 3I). In fasted animals, following an injection of 40ng/g of body weight of ucOCN, the maximum serum concentrations of ucOCN is reduced at 30 min compared to the level reached in fed animals, regardless of glycosylation status (compare Figure 3J and 3H). However, the level of glycosylated ucOCN remains higher compared to the non-glycosylated form for the following 90 minutes. These results establish that *O-*glycosylation increases the stability of mouse OCN protein in vivo.

### Human OCN is not glycosylated

Sequence alignments revealed that the residue corresponding to S8 in the mouse protein is a tyrosine (Y12) in human OCN (Figure 4A). In addition, human OCN does not contain any serine or threonine residues and migrates at a lower molecular weight compared to mouse OCN when expressed and secreted by osteoblasts, HEK293 or CHO cells (Figure 4B and data not shown). Since mouse and human ucOCN have a very similar predicted molecular weight, i.e., 5.1 and 5.8 kDa respectively, these observations suggested that human OCN may not be *O-*glycosylated. Remarkably, introduction of a single serine residue (Y12S mutation) in the human protein is sufficient to induce its *O-*glycosylation in osteoblasts as visualized by Western blot (Figure 4C). In contrast, introducing a leucine at the same position (Y12L) did not alter human OCN apparent molecular weight, indicating that this tyrosine residue is not normally subjected to *O-*glycosylation. Since the apparent molecular weight of both native and Y12S human OCN are increased following treatment with RVKR, we concluded that *O*-glycosylation does not affect human OCN processing by furin (Figure 4D). These results establish that mature human OCN is not normally subjected to *O-*glycosylation, but that a single amino acid change (Y12S) is sufficient to induce its *O*-glycosylation in osteoblasts.

**Figure 4.**
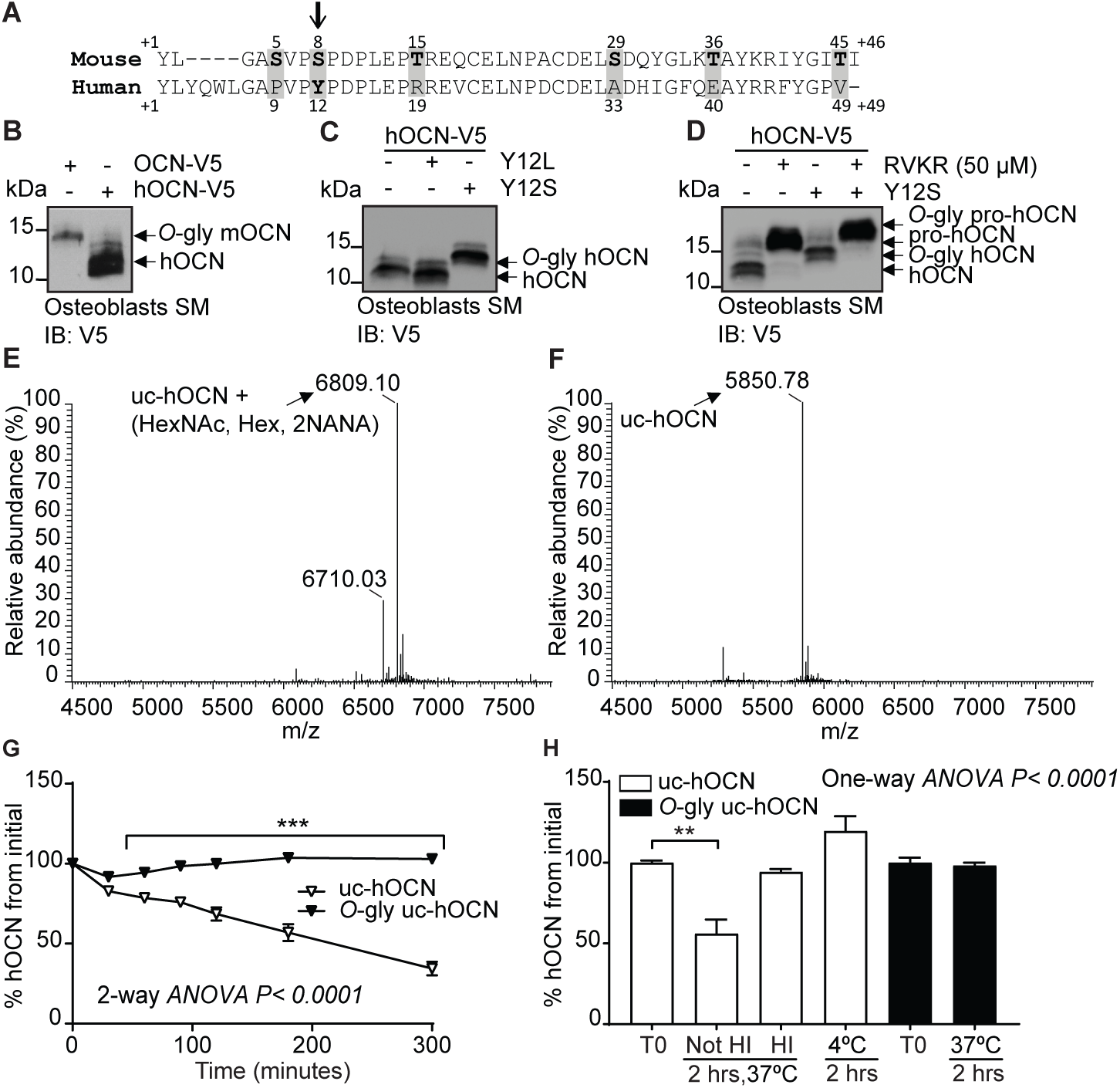
Human OCN O-glycosylation increases its half-life ex vivo. **(A)** Amino acid alignment of mouse and human OCN. The six serine and threonine residues present in the mouse protein and their corresponding amino acids in human OCN are highlighted in grey. The site of O-glycosylation in mouse OCN (S8) is indicated by an arrow. **(B)** Western Blot analysis on the secretion media (SM) of primary osteoblasts transfected with human OCN-V5 (hOCN) and mouse OCN-V5 (OCN). **(C)** Western blot analysis on the secretion media (SM) of primary osteoblasts transfected with hOCN-V5 containing or not the indicated mutations. **(D)** Western Blot analysis on the secretion media (SM) of primary osteoblasts transfected with hOCN-V5 containing or not the Y12S mutations and treated or not with 50 μM Dec-RVKR-CMK (RVKR). **(E)** Annotated and deconvoluted MS spectrum of purified O-glycosylated uncarboxylated human OCN (O-gly uc-hOCN). **(F)** Annotated and deconvoluted MS spectrum of and purified non-glycosylated uncarboxylated human OCN (uc-hOCN). **(G-H)** Ex vivo half-life of O-gly uc-hOCN and uc-hOCN in mouse plasma. **(G)** 60 ng/ml of O-gly uc-hOCN and uc-hOCN were incubated at 37°C in plasma of OCN deficient mice (Ocn-/-) (n=4) for 0 to 5 hours and hOCN levels were measured at the indicated time. **(H)** O-gly uc-hOCN and uc-hOCN were incubated in Ocn-/- plasma for 2 hours at 37°C in different conditions or at 4°C. T0: start point, see material and methods; HI: heat inactivated plasma; HexNAc: N-acetylhexosamine; Hex: Hexose; NANA: N-acetylneuraminic acid. Uc-hOCN levels were measured at the indicated time points using an uc-hOCN ELISA assay. Results are given as mean ±SEM. ** p<0.01; *** p<0.001 using 2-way ANOVA for repeated measurements with Bonferroni multiple comparisons test.

Because *O*-glycosylation impacts mouse ucOCN half-life in plasma, we next tested whether this PTM had a similar effect on human ucOCN. We produced and purified *O*-glycosylated human ucOCN^Y12S^ from HEK293 and compared its purity and molecular weight to native non-glycosylated human ucOCN by LC-MS, LC-MS/MS and SDS-PAGE (Figure 4E, 4F and Supplemental Figure 2E-F). Confirming what was observed in osteoblast, the ucOCN containing the Y12S mutation purified form HEK293 was found to be O-glycosylated (Figure 4E). These proteins were then incubated in *Ocn-/-* mouse plasma at 37°C and the concentration of ucOCN monitored over time using a specific ELISA assay (Lacombe et al., 2020). As shown in Figure 4G, in the conditions of this assay, non-glycosylated human ucOCN level declines by 50% within 180 min, while the concentration of the *O-*glycosylated version remains stable over the course of the experiment (i.e., 5 hours). As observed with the mouse protein, human ucOCN degradation was only inhibited when the plasma was heat-inactivated or incubated at 4°C (Figure 4H and Supplemental Figure 2G), suggesting that glycosylation protects mouse and human ucOCN from degradation through a similar mechanism.

## DISCUSSION

In this study we identified *O*-glycosylation as a novel PTM regulating mouse ucOCN half-life in the circulation. We also showed that *O*-glycosylation is not found on human mature OCN, but that *O*-glycosylation of human OCN by means of a single amino acid change can improve its half-life in plasma ex vivo. These findings reveal an important species difference in the regulation of OCN and may also have important implication for the future use of recombinant ucOCN as a therapeutic agent in humans.

Numerous secreted protein are subjected to mucin-type (GalNAc-type) *O*-glycosylation, a PTM which is initiated in the Golgi apparatus and involves multiple sequential glycosylation steps to produce diverse *O*-glycan structures (Steentoft et al., 2013, Bennett et al., 2012). The initiation of *O*-glycosylation is catalyzed by the GalNAc-Ts isoenzymes, however, the specific protein sequence(s) targeted by each of the GalNAc-Ts remain poorly characterized, although weak acceptor motifs for these have been identified by in vitro analyses (Perrine et al., 2009, Gerken et al., 2006). Here, we demonstrate that mouse OCN is *O*-glycosylated likely on serine 8, which is located within the amino acid sequence SVP<u>S</u>PDP^11^. Interestingly, this sequence strongly matches the consensus site previously defined for GalNAc-T1 and GalNAc-T2 (Gerken et al., 2006). In particular, the presence of proline residues in position -1, +1 and +3 has been shown to be determinant in the recognition of peptide substrates by GalNAc-T1 and GalNAc-T2 in vitro. We used an isogenic cell library with combinatorial engineering of isoenzyme families to explore the regulation of osteocalcin O-glycosylation by GalNAc-Ts (Narimatsu et al., 2019a, Narimatsu et al., 2019b). Combined knock out of GalNAc-T1, 2 and 3 only partially abolishes mouse OCN *O*-glycosylation in HEK293 cells, suggesting that OCN may be a substrate for additional GalNAc-Ts.

*O*-glycosylation was shown to interfere with the action of proprotein convertases on several pro-hormones and receptors (Goth et al., 2015, Kato et al., 2006, May et al., 2003, Schjoldager and Clausen, 2012, Schjoldager et al., 2010, Goth et al., 2017). This appears not to be the case for OCN as its *O*-glycosylation does not interfere with its processing by furin in vitro and in vivo. In other proteins, such as leptin and erythropoietin, glycosylation adducts were shown to increase protein stability and half-life in circulation (Elliott et al., 2003, Creus et al., 2001). More recently, O-glycosylation was shown to protect atrial natriuretic peptide from proteolytic degradation by neprilysin or insulin-degrading enzyme in vitro (Hansen et al., 2019). Additional studies showed that sialic acid residues present in the glycosylation adducts increase protein charge, thereby improving serum half-life and decreasing liver and renal clearance (Morell et al., 1971, Runkel et al., 1998, Perlman et al., 2003, Ziltener et al., 1994). Although our data suggest that protection from proteolytic cleavage by plasmin in the plasma might be one mechanism by which *O-*glycosylation extend OCN half-life, we cannot exclude that in vivo, *O-*glycosylation may also decrease liver and renal clearance of OCN.

Mouse OCN contains arginine (R) residues at position 16 and 40, corresponding to R20 and R44 in human OCN, which could be potential cleavage sites for plasmin (Figure 4A). A putative plasmin cleavage site is present in mouse (i.e., AYK↓R^40^, where the arrow indicates the cleavage site) and human (i.e., AYR↓R^44^) OCN at the corresponding position (Backes et al., 2000). Bovine OCN, like the human protein, does not possess the *O-*glycosylation site found in mouse, and was shown to be cleaved by plasmin in vitro between R43 and R44 (Novak et al., 1997). Here we show that plasmin is sufficient to reduce the concentration non-glycosylated ucOCN, by more than 90% in two hours, confirming that mouse ucOCN is also a plasmin substrate. Supporting a role for plasmin in the cleavage of mouse OCN in vivo, OCN serum levels are decreased in mice deficient in plasmin activator inhibitor I (PAI-I), which have higher plasmin activity (Tamura et al., 2013). Interestingly, circulating plasmin activity increases with age in rats and humans (Paczek et al., 2009, Paczek et al., 2008), an observation which could in part explain the gradual reduction of serum OCN concentrations during aging (Mera et al., 2016a). Notably, a nonsynonymous rare variant (rs34702397) in OCN resulting in the conversion of R43 in a glutamine (Q) exists in humans of African ancestry and was nominally associated with insulin sensitivity index and glucose disposal in a small cohort of African Americans (Das et al., 2010). Since the R43Q variant eliminates the potential plasmin cleavage site in the C-terminal region, it will be interesting to further investigate if subjects carrying this variant have higher OCN circulating levels.

We found that human OCN is not subjected to *O*-glycosylation and that consequently it has a reduced half-life in plasma ex vivo. The *O*-glycosylation sequence “SVP<u>S</u>PDP^11^” of

mouse OCN is conserved in the human protein, except for the amino acids corresponding to S5 and S8, which are replaced by a proline (P9) and a tyrosine (Y12) (i.e., “PVP<u>Y</u>PDP^15^”). Remarkably, we could introduce *O*-glycosylation into human OCN by a single amino acid change (Y12S). *O*-glycosylated human OCN is protected from degradation in plasma ex vivo similarly to the glycosylated mouse protein. Hence, this difference in the *O-*glycosylation status of OCN could potentially explain why circulating level of OCN in 1- to 6-month-old mice is 5-10 times higher than the level measured in young or adult human (Supplemental table 1).

It remains unknown if the increased half-life of *O*-glycosylated ucOCN will result in improved biological activity in vivo, although it was shown to be the case for other proteins (Baudys et al., 1995, Runkel et al., 1998, Elliott et al., 2003). Moreover, even though it is currently not known if *O-*glycosylated ucOCN is active in mice, a few evidences suggest that ucOCN *O-*glycosylation does not interfere with its activity. First, our data show that mouse OCN is endogenously *O*-glycosylated in vivo and that more than 80% of OCN, including the ucOCN fraction, is *O*-glycosylated in osteoblast supernatant and in serum. This suggest that the bioactive form of OCN is *O-*glycosylated in vivo in mice. Second, bacterially produced mouse ucOCN, which is not *O*-glycosylated, was previously shown to be bioactive in cell based assays and in vivo (Pi et al., 2011, Zhou et al., 2013, Gupte et al., 2014) suggesting that *O*-glycosylation is not required for ucOCN activity in mice.

In summary, this work identified *O*-glycosylation as a previously unrecognized OCN PTM regulating its half-life in circulation in mice. This modification is not conserved in human yet introducing *O-*glycosylation in human ucOCN also increases its half-life in plasma. These findings reveal an important difference between mouse and human OCN biology and also provide an approach to increase recombinant human OCN half-life in vivo, which might be relevant for the future development of OCN-based therapies for human diseases.

## MATERIAL AND METHODS

### Animal models

The *Furin*^*fl/fl*^ and *Furin*^*osb*^*-/-* mice were generated by breeding *Furin*^*fl/fl*^ with *OCN-Cre* transgenic mice that express Cre recombinase under the control of human OCN promoter as described previously (Al Rifai et al., 2017). *Ocn-/-* mice were generated using homologous recombination to replace *Ocn1* (*Bglap1*) and *Ocn2* (*Bglap2*) genes in the mouse *Ocn* cluster with a neomycin resistance cassette (Ducy et al., 1996). All strains used in this study were backcrossed on a C57BL/6J genetic background more than 10 times and maintained under 12-hour dark/12-hour light cycles in a specific pathogen–free animal facility (SPF) at IRCM. Male mice were used in all experiments, and they were fed a normal chow diet. All animal use complied with the guidelines of the Canadian Committee for Animal Protection and was approved by IRCM Animal Care Committee.

### DNA constructs

Mouse pro-OCN cDNA was cloned into the pIRES2-EGFP-V5 plasmid in EcoRI and AgeI cloning sites. S5A/S8/AT15A pro-OCN, S29A/T36A/T45A pro-OCN and S5S/S8A/T15A/S29A/T36A/T45A (i.e., 6XST→6XA) pro-OCN mutant were purchased from Thermo Fisher. Human pre-pro-OCN cDNA cloned into pcDNA3 was purchased from GenScript. Each construct was used as PCR template for amplification and to introduce EcoRI and AgeI cloning sites and cloned in pIRES2-EGFP-V5 plasmid. Point mutations in mouse pro-OCN (S5A, S8A, T15A) and Y12S in human pro-OCN were generated by site directed mutagenesis using specific primer (Supplemental Table 5).

The cDNA coding of the Fc and hinge region of human immunoglobulin flanked with HindIII-BamHI restriction sites was amplified using standard PCR and pTT5-Fc1_CTL vector as a template (Saavedra et al., 2013). The PCR product was cloned in pcDNA3.1-myc-His B in HindIII-BamHI cloning site, generating the pcDNA3.1-Fc-hinge-myc-His vector. The cDNA coding for mature hOCN^(Y12S)^ was generated using pIRES2-EGFP-hOCN (Y12S)-V5 as a template, to which a thrombin (Thr) cleavage site was added at the N-terminus and BglII and EcoRI restriction sites were introduced by standard PCR amplifications. The Thr-hOCN^(Y12S)^ product was cloned in the pcDNA3.1-Fc-hinge-myc-His vector. The generated vector pcDNA3.1-Fc-hinge-Thr-hOCN^(Y12S)^ is an expression vector of human OCN fusion protein composed of the Fc and hinge region of human IgG1, thrombin cleavage site and human OCN (Y12S). Mouse OCN fused to Fc were generated following the same procedure and using different primers.

### Cell culture and transfection

Primary osteoblasts were prepared following a previously described protocol (Ferron et al., 2015). In brief, calvariae were collected from 3 days old mice and washed with 1× PBS and digested 2 times for 10 minutes in digestion solution (αMEM, 0.1 mg/ml collagenase type 2 [Worthington Biochemical Corporation] and 0.00075% trypsin) that was discarded after incubation. Following two 30 minutes incubations, the digestion solutions were collected, centrifuged and cells recovered were cultured in αMEM supplemented with 10% FBS, penicillin and streptomycin (PS), and L-glutamine. Culture media was supplemented with 5 mM β-glycerophosphate and 100 µg/ml L-ascorbic acid to induce osteoblasts differentiation and it was replaced every 2 days for 21 days.

Primary osteoblasts were transfected using jetPRIME Reagent (Polypus transfection). After an overnight incubation, media were changed to secretion media (FBS-free αMEM plus 2mM L-glutamine, PS). After 24 hours of secretion, media were collected, and cells were lysed in protein lysis buffer (20 mM Tris-HC pH 7.4, 150 mM NaCl, 1 mM EDTA, 1 mM EGTA, 1% Triton, 1 mM PMSF, and 1× protease inhibitor cocktail) and analyzed by Western blotting. In some experiments, osteoblasts were treated with the γ-carboxylation inhibitor warfarin (50 μM; Santa Cruz Biotechnology), the *N-*acetylgalactosaminyltransferase inhibitor GalNAc-bn (2 mM, Sigma) or the proprotein convertase inhibitor Dec-RVKR-CMK (50 μM, Tocris), combined with 22 μM vitamin K_1_ (Sigma).

Chinese hamster ovary (CHO) cells, originally purchased from ATCC, and Chinese hamster ovary ldlD cells (CHO-ldlD; originating from the M. Krieger laboratory (Kingsley et al., 1986)) were cultured in DMEM-F12 containing PS and 5% FBS for CHO cells or 3% FBS for CHO-ldlD cells and transfected using Lipofectamine 2000 (Thermo Fisher) following standard protocol. Secretion was performed in DMEM-F12 media supplemented with PS and 22 µM VK_1_. In some experiments, CHO-ldlD culture, transfection and secretion media was supplemented with 0.1 mM galactose and/or 1 mM *N*-acetylgalactosamine (GalNAc) to rescue the *O-*glycosylation defect as previously reported (Kingsley et al., 1986)

Human embryonic kidney cells HEK293 were originally purchased from ATCC. *COSMC* knockout HEK293sc (HEK293 simple cell) cells and *GALNTs* deficient HEK293 cells were generated using Zinc finger nuclease (ZFN) gene editing as described previously (Goth et al., 2015, Schjoldager et al., 2012, Steentoft et al., 2013, Steentoft et al., 2011, Goth et al., 2017). Cells were transfected using Lipofectamine 2000 reagent and secretion was performed over 24 hours in EMEM supplemented with PS and 22 µM VK1. In some experiments, HEK293 cells were treated with warfarin, GalNAc-bn or Dec-RVKR-CMK combined with 22 μM vitamin K_1_.

For Western blot analysis, proteins were resolved on 15% Tris-tricine gel and blotted overnight with indicated antibody. Antibody used in this study are: anti-V5 (mouse, clone V5-10, V8012; Sigma-Aldrich), anti–β-actin (mouse, clone AC-15, A5441; Sigma-Aldrich), anti-GFP (mouse, clones 7.1 and 13.1, 11814460001; Sigma), anti-Gla OCN goat antibody which recognize amino acids 11-26 of carboxylated mature OCN and anti-Cterm OCN goat antibody which recognize amino acids 26-46 of mature mouse OCN (Ferron et al., 2010b).

### In vitro de-glycosylation assay

Flushed mouse femur and tibia from C57BL/6J were homogenized in lysis buffer (20 mM Tris-HCl pH 7.4, 150 mM NaCl, 1 mM EDTA, 1 mM EGTA, 1% Triton, 1 mM PMSF, and 1× protease inhibitors cocktail). Tissue homogenates were then centrifuged for 10 minutes at 4000 rpm to remove insoluble material. In vitro de-glycosylation assay was performed on 10 μg of bone homogenate. Briefly, proteins were denatured in denaturing buffer (0.5% SDS, 40 mM DTT) at 95°C for 5 min and incubated with 80000 units of *O*-glycosidase and 100 units of neuraminidases for 4 hours at 37°C following the NEB kit protocol (E0540S; NEB). Samples were resolved on 15% Tris-tricine SDS-PAGE gel and blotted using anti-Cterm OCN goat antibody.

### Top-down LC-MS/MS analysis MS analysis of OCN in osteoblasts secretion media and bone extracts

Differentiated osteoblasts secretion media was spun down at 1500 rpm for 5 minutes to remove cells debris. The supernatant was then incubated overnight at 4°C with anti-Cterm OCN antibody in the presence of 1× protease inhibitors cocktail. For bone extract, flushed femur and tibia from wild type mice were homogenized in lysis buffer containing (20 mM Tris-HCl pH 7.4, 150 mM NaCl, 1 mM EDTA, 1 mM EGTA, 1% Triton, 1 mM PMSF, and 1× protease inhibitors cocktail). 100 μg of protein homogenate was diluted in 1,6 ml of 100 mM phosphate buffer pH 7.4 and incubated overnight at 4°C with anti-OCN antibody. After overnight incubation, samples were centrifuged at 10000 rpm for 10 minutes to remove precipitate and supernatant was incubated with protein-G agarose beads pre-washed with 1X PBS. After for 4 hours of rotation at 4°C, beads were spun down, washed twice with 1X PBS and three times with 50 mM Ammonium Bicarbonate pH 8.0. OCN was then eluted with 100 μl of 0.5 M NH_4_OH, snap frozen in liquid nitrogen and evaporated under vacuum using speedvac concentrator (Thermo scientific).

Samples were diluted in 25% ACN 0.3%TFA and loaded onto a 50×4.6 mm PLRP-S 300A column (Agilent Technologies) connected to an Accela pump (Thermo Scientific) and a RTC autosampler (Pal systems). The buffers used for chromatography were 0.1% formic acid (buffer A) and 100% acetonitrile/0.1% formic acid (buffer B). Proteins and peptides were eluted with a two slopes gradient at a flowrate of 120 µL/min. Solvent B first increased from 12 to 50% in 4.5 min and then from 50 to 70% in 1.5 min. The HPLC system was coupled to a Q Exactive mass spectrometer and an Orbitrap Fusion (Thermo Scientific) through an electrospray Ion Source. The spray and S-lens voltages were set to 3.6 kV and 60 V, respectively. Capillary temperature was set to 225 °C. Full scan MS survey spectra (m/z 600-2000) in profile mode were acquired in the Orbitrap with a resolution of 70,000 or 120,000 with a target value at 3e6. The 4 most intense protein/peptide ions were fragmented in the HCD (higher-energy collision dissociation) collision cell and analyzed in the Orbitrap with a target value at 5e5 and a normalized collision energy at 33. Data processing protocol: the identification of the different forms of OCN was performed by manual denovo sequencing.

### *Galnts* expression in osteoblasts

RNA was extracted from non-differentiated and differentiated calvariae osteoblasts using Trizol reagent (Thermo Fisher Scientific) following standard protocol. RNA was treated with DNAse I and reverse transcribed using poly dT primers, random primers and MMLV reverse transcriptase (Thermo Fisher Scientific). QPCR was performed on standards of diluted genomic DNA and cDNA products using specific primers (Supplemental table 5) on a ViiA 7 Real-Time PCR system (Thermo Fisher Scientific). *Galnts* gene copy numbers were calculated using the genomic DNA as a standard curve and variation between biological replicate was normalized using *Actb* expression level.

### Generation of stable HEK293 clones expressing mouse and human OCN fused to the Fc region of human immunoglobulin

To generate stable clonal cell lines expressing glycosylated human and mouse OCN, HEK293 were transfected with pcDNA3.1-Fc-hinge-Thr-hOCN^Y12S^ and pcDNA3.1-Fc-hinge-Thr-OCN respectively using Lipofectamine 2000 reagent. Following 48 hours of transfection, cells were trypsinized and resuspended in sorting buffer containing (1× sterile PBS, 2% FBS and 1 mM EDTA). Cells were sorted at a concentration of 5-10 cells/well in 96 well plates containing the selection media (EMEM, 10% FBS supplemented with G418 sulfate (500 μg/ml; Wisent). Following two weeks of selection, isolated colonies appeared and the expression of mouse and human OCN was assessed using ELISA assay described below. Clones expressing the highest levels of OCN were amplified and frozen.

### Purification of mouse and human OCN fused to the Fc region of human immunoglobulin

TM102F12 clone expressing IgFc-mOCN fusion protein and 22H5 clone expressing IgFc-hOCN^Y12S^ fusion protein were cultured in triple layer 175cm^2^ flasks. After reaching 100% confluency, cells were kept in secretion media (EMEM media supplemented with 1% FBS and 10 μM warfarin to block γ-carboxylation) for 72 hours. Secretion media was collected, filtered with 0.45 μm filter, and media was pH buffered with 10× binding buffer (0.2 M phosphate buffer, pH 7). This cell supernatant was then loaded into protein A affinity column (HiTrap protein A high performance, GE29-0485-76; GE Healthcare Life Sciences,) using liquid chromatography system (GE AKTA Prime Plus). Column was then washed with 20 ml of 1× binding buffer (0.02 M phosphate buffer, pH 7) and 5 ml of filtered 1× PBS. To release OCN from the column, OCN fusion protein was digested with thrombin (27-0846-01, GE Healthcare Life Sciences) and eluted with 1× PBS. Thrombin was subsequently removed using benzamidine sepharose (17-5123-10, GE healthcare). Mouse and human OCN purity were assessed using Coomassie staining and liquid chromatography-mass spectrometry (LC-MS) analysis compared to purified non-glycosylated mouse or human ucOCN. Mouse OCN was quantified using ELISA assay as described previously (Ferron et al., 2010b). Human ucOCN measurements were performed using a commercially available human ucOCN ELISA (BioLegend, 446707) (Lacombe et al., 2020).

### Ex vivo half-life and plasmin enzymatic assays

The mouse OCN ex vivo half-life assays were performed with fresh plasma (lithium heparin) collected from four independent *Ocn-/-* mice. Glycosylated OCN produced in HEK293 cells and non-glycosylated OCN, produced in bacteria as previously described (Ferron et al., 2012), were incubated at 100 ng/ml in plasma at 37°C and OCN level was measured at indicated time points using the total mouse OCN ELISA assay described previously (Ferron et al., 2010b). Human OCN half-life assay was performed ex vivo using *Ocn-/-* mice plasma and human OCN at 60 ng/ml. Human OCN levels was measured at different time point using the human ucOCN ELISA described above. At the experiment start point (T0), an aliquot of plasma was diluted in ELISA assay buffer and kept on ice for OCN measurements. In some experiments, plasma was heat-inactivated for 30 minutes at 56°C, or treated with EDTA (10 mM, Wisent), phenylmethylsulfonyl fluoride (PMSF, 10 mM, Sigma), pepstatin A (Pep A, 10 μM, Sigma) which inhibits aspartic proteases (pepsin, cathepsin D, renin, chymosin), RVKR (Dec-RVKR-CMK, 50 μM; Tocris) or talabostat (10 mM; Tocris). Results are calculated in percentage relative to the initial concentration of non-glycosylated and glycosylated OCN respectively.

The plasmin enzymatic assay was performed on plasma ex vivo in the presence of non-glycosylated OCN and glycosylated OCN as follow: plasma from Ocn-/- mice was heat-inactivated for 30 minutes at 56°C, then diluted two times in the enzyme assay buffer containing 100mM Tris buffer pH 7.5. Non-glycosylated OCN or glycosylated OCN was incubated at 50ng/ml in the diluted plasma for two hours in the presence of different concentration of human plasmin (Sigma) ranging from 0 to 0.3U/ml. After two hours, OCN level was measured using total mouse OCN ELISA. Results are represented as percentage of initial concentration of non-glycosylated and glycosylated OCN respectively.

### In vivo half-life assay

For in vivo half-life assay, *Ocn-/-* male mice were injected intraperitoneally with 40 or 80 ng/g of body weight of mouse *O*-glycosylated ucOCN or non-glycosylated ucOCN, serum OCN level was analyzed at indicated time points using total mouse OCN ELISA. In all ex vivo and in vivo studies, mouse or human proteins were prepared in saline solution (0.9% NaCl) containing BSA (35 μg/ml) as a carrier.

### Statistics

Statistical analyses were performed using GraphPad Prism software version 7.03. Results are shown as the mean ± SEM. For single measurements, an unpaired, 2-tailed Student’s *t* test was used, while 1-way ANOVA followed by Bonferroni’s post-test was used for comparison of more than 2 groups. For repeated measurements (e.g., half-life study ex vivo and in vivo), a repeated-measurement 2-way ANOVA followed by Bonferroni’s post-test was used. A *P* value of less than 0.05 was considered statistically significant. All experiments were repeated at least 3 times or performed on at least 3 independent animals.

## ACKNOWLEDGEMENTS

We thank Dr. Nabil Seidah for providing reagents as well as the CHO and CHO-ldld cells. We thank Dr. John Creemers for providing the *Furin* floxed mice. This work was supported by funding from the Canada Research Chair program (M.F.), the Canadian Institutes of Health Research (M.F., MOP-133652 and PJT-159534) and the Natural Sciences and Engineering Research Council of Canada (RGPIN-2016-05213, to MF), and the Danish National Research Foundation (DNRF107, to HC). OAR received scholarships from IRCM and FRQS.

---

**Supplemental figure 1.**
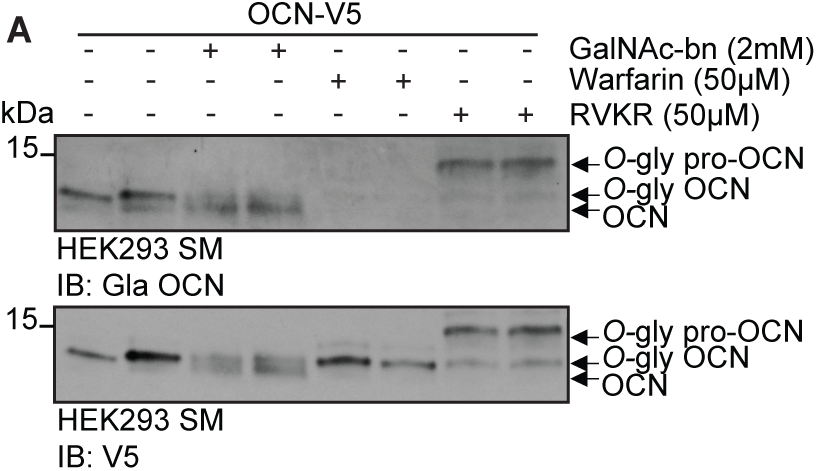
Mouse OCN O-glycosylation occurs independently of its carboxylation and processing in HEK293 cells. **A)** Western blot analysis on the secretion media (SM) of HEK293 cells transfected with mouse OCN-V5 and treated or not with 2 mM of GalNAc-bn, 50 μM warfarin or 50 μM Dec-RVKR-CMK (RVKR).

**Supplemental figure 2.**
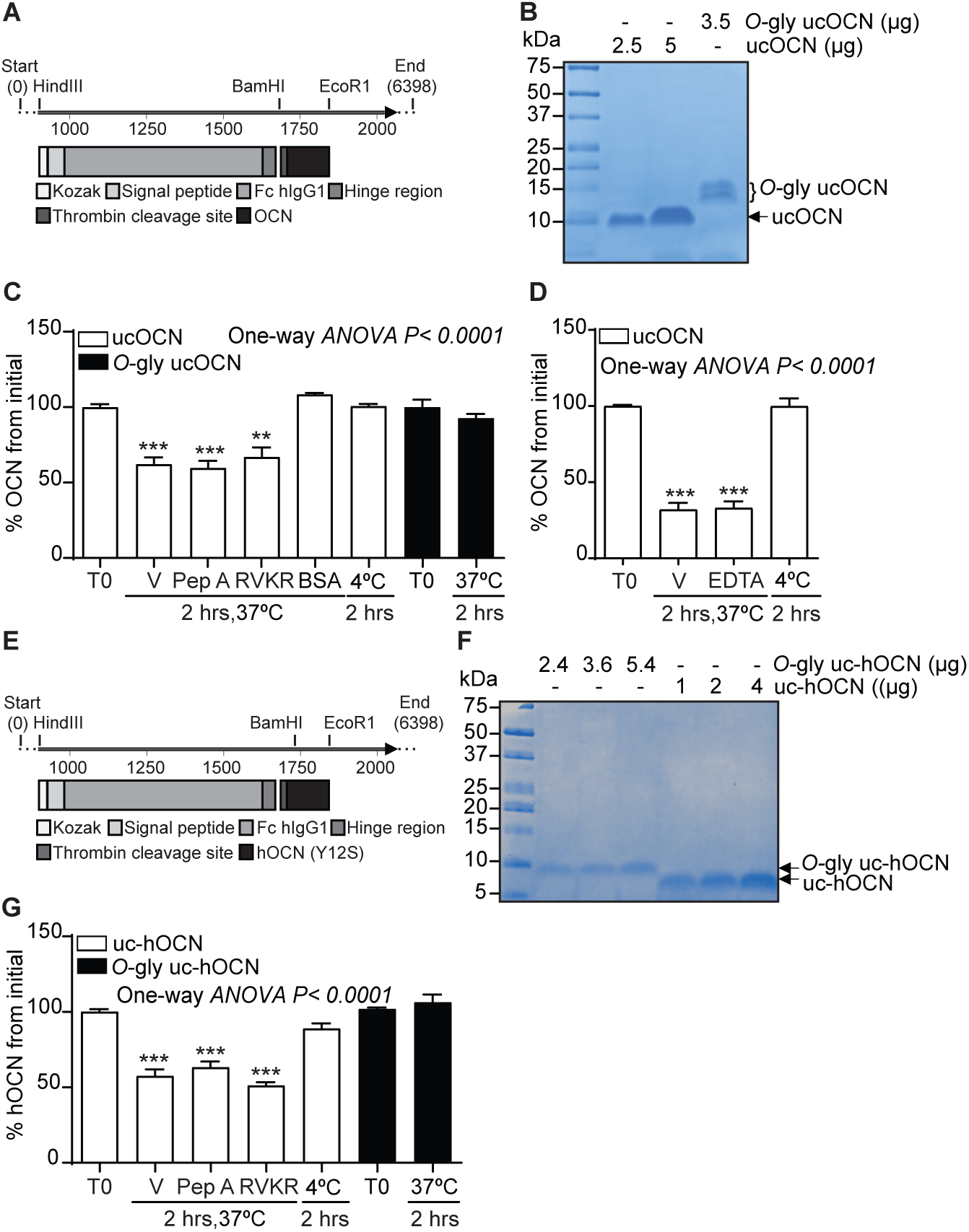
Purification of recombinant O-glycosylated mouse and human ucOCN and effect of different protease inhibitors on non-glycosylated mouse and human ucOCN plasma half-life. **(A)** Map of the pcDNA3.1-Fc-hinge-Thr-OCN construct used to produce and purify mouse ucOCN fusion protein. **(B)** Coomassie staining of purified O-glycosylated mouse ucOCN (O-gly ucOCN) compared to non-glycosylated mouse ucOCN (ucOCN) produced in bacteria. **(C-D)** Ex vivo half-life of O-gly ucOCN and ucOCN in Ocn-deficient plasma (n=3-4 plasma). **(C)** 100 ng/ml O-gly ucOCN and ucOCN were incubated for 2 hours in plasma at 37°C or 4°C and treated with vehicle (V) or with protease inhibitors or incubated in 3.5% BSA (bovine serum albumin prepared in saline solution). **(D)** 100 ng/ml O-gly ucOCN and ucOCN was incubated for 2 hours in normal plasma at 37°C or 4°C and treated with vehicle (V) or with EDTA (10 mM). **(E)** Map of pcDNA3.1-Fc-hinge-Thr-hOCN (Y12S) construct used to produce and purify O-glycosylated human ucOCN fusion protein. **(F)** Coomassie staining of purified O-glycosylated human ucOCN (O-gly uchOCN) compared to non-glycosylated human ucOCN (uc-hOCN) produced in bacteria. **(G)** Ex vivo half-life of O-gly uc-hOCN and uc-hOCN in Ocn deficient plasma (n=3-4 plasma). 60 ng/ml O-gly uc-hOCN and uc-hOCN were incubated for 2 hours in normal plasma at 37°C or 4°C and treated with vehicle (V) or with protease inhibitors. T0: start point, see material and methods; Pep A: 10 μM Pepstatin A; RVKR: 50 μM Dec-RVKR-CMK.

**Supplemental table 1:** OCN serum levels in mouse and human at different ages.

| OCN serum levels (ng/ml) |  |  |
| --- | --- | --- |
| Age (mice/human) | Mouse [mean ± SD (n)] | Human* [mean ± SD (n)] |
| 2 weeks/ 1 year old | 1369.7 ± 146.69 (8) | 62.9 ± 8.14 (43) |
| 4 weeks/ 11-13 years old | 617.2 ± 192.50 (5) | 74.1 ± 8.88 (41) |
| 13 weeks/ 25-29 years old | 252.2 ± 7.96 (4) | 21 ± 6.29 (49) |
| 60 weeks/ 50-54 years old | 50 ± 7.18 (4) | 13.5 ± 6.30 (127) |

\*(Hannemann et al., 2013, Cioffi et al., 1997)

**Supplemental table 2:**
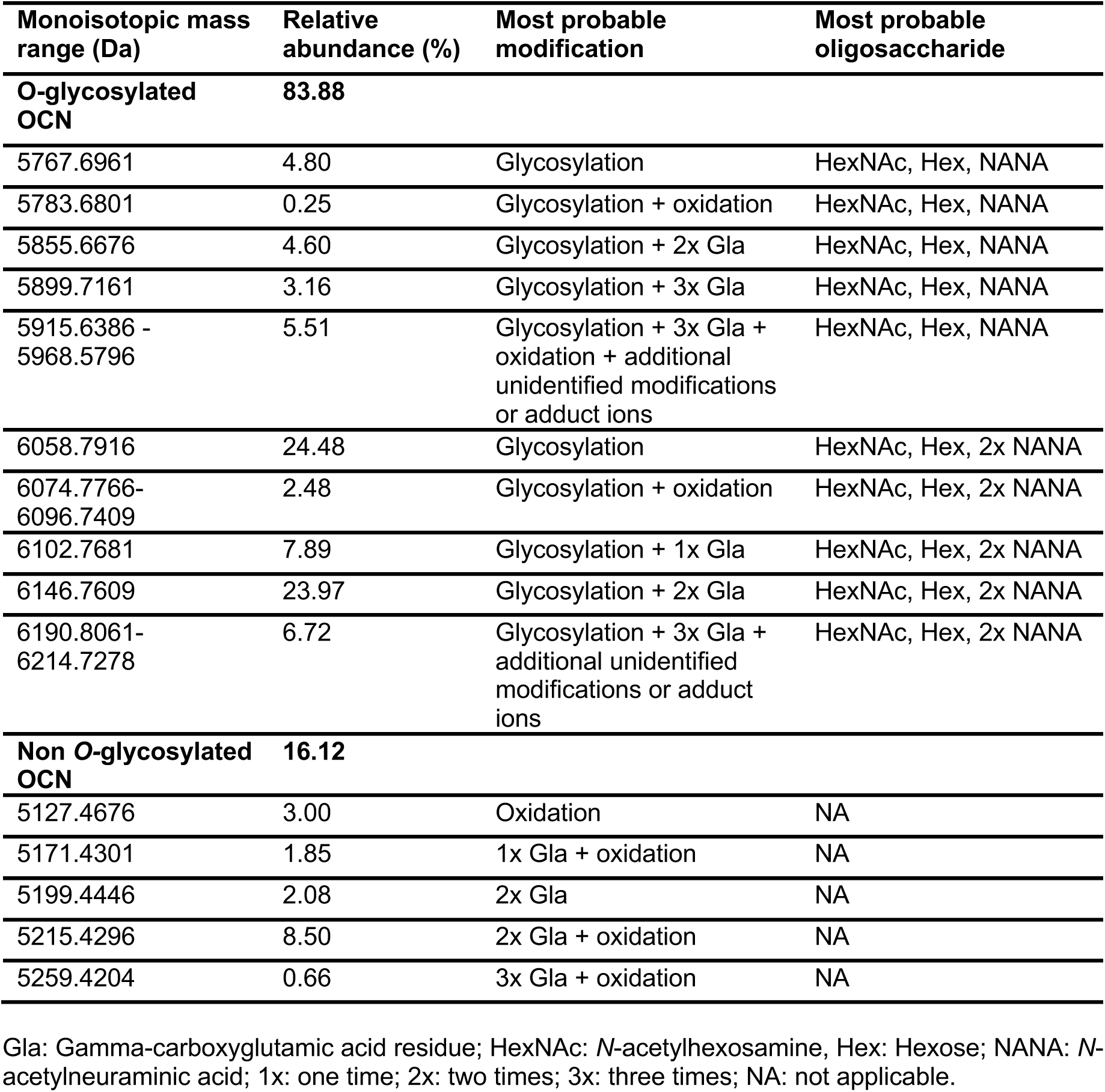
The monoisotopic mass and relative abundance of the different OCN forms detected in the supernatant of differentiated osteoblasts.

**Supplemental table 3:** The monoisotopic mass and relative abundance of the different OCN forms detected in mouse bone homogenates.

| **Monoisotopic mass range** | **Relative abundance (%)** | **Most probable modification** | **Most probable oligosaccharide** |
| --- | --- | --- | --- |
| **O-glycosylated OCN** | **99.07** |  |  |
| 5855.6676 | 0.54 | Glycosylation + 2x Gla | HexNAc, Hex, NANA |
| 5899.7161 | 7.43 | Glycosylation + 3x Gla | HexNAc, Hex, NANA |
| 5915.6386-6135.6796 | 36.07 | Glycosylation + 3x Gla + oxidation + additional unidentified modifications or adduct ions | HexNAc, Hex, NANA |
| 6146.7609-6162.7991 | 5.42 | Glycosylation + 2x Gla + additional unidentified modifications | HexNAc, Hex, 2xNANA |
| 6190.8061 | 4.49 | Glycosylation + 3x Gla | HexNAc, Hex, 2xNANA |
| 6206.8016-6441.7636 | 45.12 | Glycosylation + 3x Gla + oxidation + additional unidentified modifications or adduct ions | HexNAc, Hex, 2xNANA |
| **Non O-glycosylated OCN** | **0.93** |  |  |
| 5259.4204 | 0.93 | 3x Gla + oxidation | NA |

Gla: Gamma-carboxyglutamic acid residue; HexNAc: *N*-acetylhexosamine, Hex: Hexose; NANA: *N*-acetylneuraminic acid; 1x: one time; 2x: two times; 3x: three times; NA: not applicable.

**Supplemental table 4:**
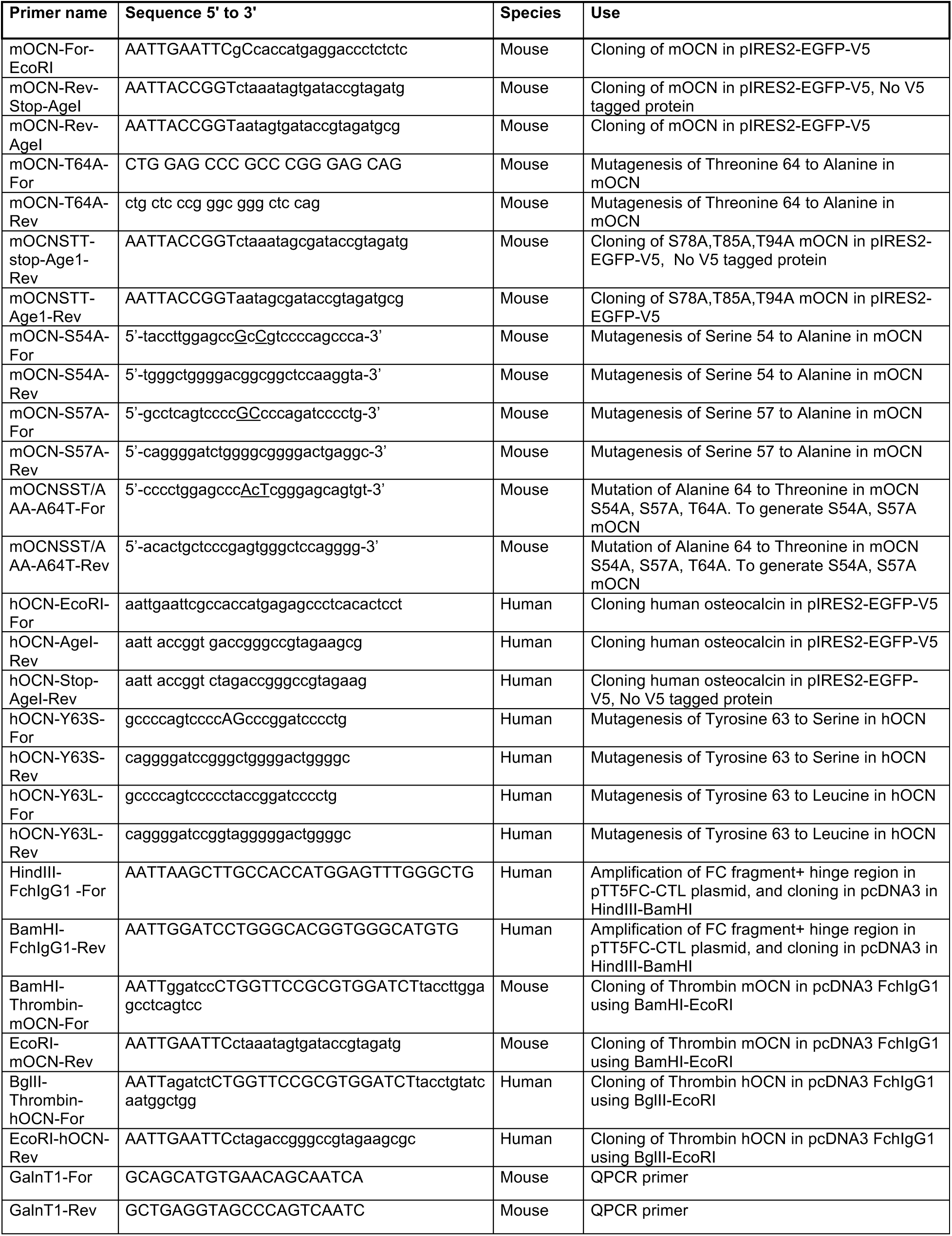
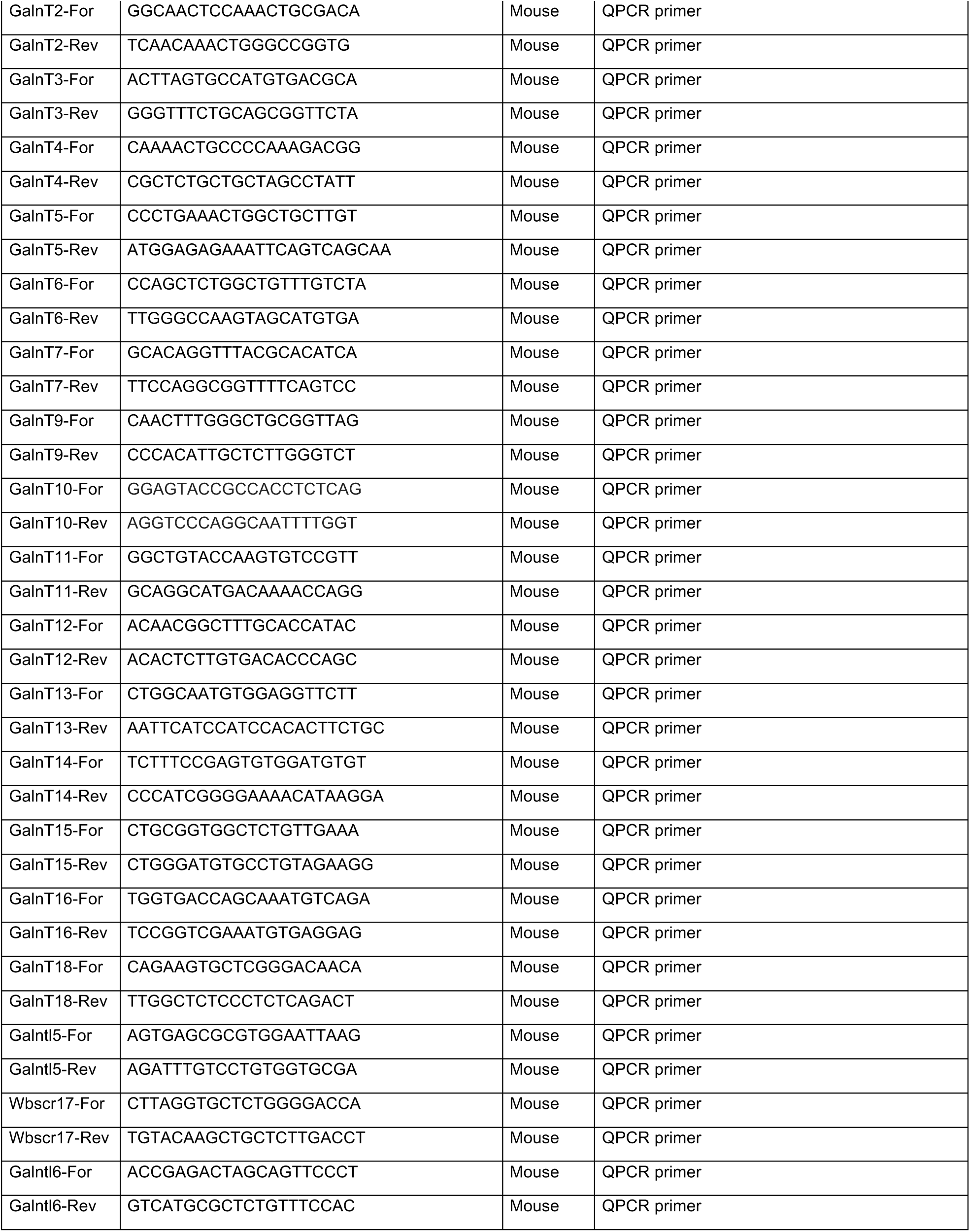
Sequence of oligonucleotides used.

## REFERENCES

Al Rifai, O., Chow, J., Lacombe, J., Julien, C., Faubert, D., Susan-Resiga, D., Essalmani, R., Creemers, J. W., Seidah, N. G. & Ferron, M. 2017. Proprotein convertase furin regulates osteocalcin and bone endocrine function. J Clin Invest, 127, 4104–4117.

Backes, B. J., Harris, J. L., Leonetti, F., Craik, C. S. & Ellman, J. A. 2000. Synthesis of positional-scanning libraries of fluorogenic peptide substrates to define the extended substrate specificity of plasmin and thrombin. Nat Biotechnol, 18, 187–93.

Banbula, A., Bugno, M., Goldstein, J., Yen, J., Nelson, D., Travis, J. & Potempa, J. 2000. Emerging family of proline-specific peptidases of Porphyromonas gingivalis: purification and characterization of serine dipeptidyl peptidase, a structural and functional homologue of mammalian prolyl dipeptidyl peptidase IV. Infect Immun, 68, 1176–82.

Baudys, M., Uchio, T., Mix, D., Wilson, D. & Kim, S. W. 1995. Physical stabilization of insulin by glycosylation. J Pharm Sci, 84, 28–33.

Bennett, E. P., Mandel, U., Clausen, H., Gerken, T. A., Fritz, T. A. & Tabak, L. A. 2012. Control of mucin-type O-glycosylation: a classification of the polypeptide GalNAc-transferase gene family. Glycobiology, 22, 736–56.

Berger, J. M., Singh, P., Khrimian, L., Morgan, D. A., Chowdhury, S., Arteaga-Solis, E., Horvath, T. L., Domingos, A. I., Marsland, A. L., Yadav, V. K., Rahmouni, K., Gao, X. B. & Karsenty, G. 2019. Mediation of the Acute Stress Response by the Skeleton. Cell Metab, 30, 890–902 e8.

Bermpohl, F., Loster, K., Reutter, W. & Baum, O. 1998. Rat dipeptidyl peptidase IV (DPP IV) exhibits endopeptidase activity with specificity for denatured fibrillar collagens. FEBS Lett, 428, 152–6.

Cioffi, M., Molinari, A. M., Gazzerro, P., Di Finizio, B., Fratta, M., Deufemia, A. & Puca, G. A. 1997. Serum osteocalcin in 1634 healthy children. Clin Chem, 43, 543–5.

Coutts, S. J., Kelly, T. A., Snow, R. J., Kennedy, C. A., Barton, R. W., Adams, J., Krolikowski, D. A., Freeman, D. M., Campbell, S. J., Ksiazek, J. F. & Bachovchin, W. W. 1996. Structure-activity relationships of boronic acid inhibitors of dipeptidyl peptidase IV. 1. Variation of the P2 position of Xaa-boroPro dipeptides. J Med Chem, 39, 2087–94.

Creus, S., Chaia, Z., Pellizzari, E. H., Cigorraga, S. B., Ulloa-Aguirre, A. & Campo, S. 2001. Human FSH isoforms: carbohydrate complexity as determinant of in-vitro bioactivity. Mol Cell Endocrinol, 174, 41–9.

Das, S. K., Sharma, N. K. & Elbein, S. C. 2010. Analysis of osteocalcin as a candidate gene for type 2 diabetes (T2D) and intermediate traits in Caucasians and African Americans. Dis Markers, 28, 281–6.

De Toni, L., Guidolin, D., De Filippis, V., Tescari, S., Strapazzon, G., Santa Rocca, M., Ferlin, A., Plebani, M. & Foresta, C. 2016. Osteocalcin and Sex Hormone Binding Globulin Compete on a Specific Binding Site of GPRC6A. Endocrinology, 157, 4473–4486.

Di Nisio, A., Rocca, M. S., Fadini, G. P., De Toni, L., Marcuzzo, G., Marescotti, M. C., Sanna, M., Plebani, M., Vettor, R., Avogaro, A. & Foresta, C. 2017. The rs2274911 polymorphism in GPRC6A gene is associated with insulin resistance in normal weight and obese subjects. Clin Endocrinol (Oxf), 86, 185–191.

Ducy, P., Desbois, C., Boyce, B., Pinero, G., Story, B., Dunstan, C., Smith, E., Bonadio, J., Goldstein, S., Gundberg, C., Bradley, A. & Karsenty, G. 1996. Increased bone formation in osteocalcin-deficient mice. Nature, 382, 448–52.

Elliott, S., Lorenzini, T., Asher, S., Aoki, K., Brankow, D., Buck, L., Busse, L., Chang, D., Fuller, J., Grant, J., Hernday, N., Hokum, M., Hu, S., Knudten, A., Levin, N., Komorowski, R., Martin, F., Navarro, R., Osslund, T., Rogers, G., Rogers, N., Trail, G. & Egrie, J. 2003. Enhancement of therapeutic protein in vivo activities through glycoengineering. Nat Biotechnol, 21, 414–21.

Ferron, M., Lacombe, J., Germain, A., Oury, F. & Karsenty, G. 2015. GGCX and VKORC1 inhibit osteocalcin endocrine functions. J Cell Biol, 208, 761–76.

Ferron, M., Mckee, M. D., Levine, R. L., Ducy, P. & Karsenty, G. 2012. Intermittent injections of osteocalcin improve glucose metabolism and prevent type 2 diabetes in mice. Bone, 50, 568–75.

Ferron, M., Wei, J., Yoshizawa, T., Del Fattore, A., Depinho, R. A., Teti, A., Ducy, P. & Karsenty, G. 2010a. Insulin signaling in osteoblasts integrates bone remodeling and energy metabolism. Cell, 142, 296–308.

Ferron, M., Wei, J., Yoshizawa, T., Ducy, P. & Karsenty, G. 2010b. An ELISA-based method to quantify osteocalcin carboxylation in mice. Biochem Biophys Res Commun, 397, 691–6.

Gerken, T. A., Raman, J., Fritz, T. A. & Jamison, O. 2006. Identification of common and unique peptide substrate preferences for the UDP-GalNAc:polypeptide alpha-N-acetylgalactosaminyltransferases T1 and T2 derived from oriented random peptide substrates. J Biol Chem, 281, 32403–16.

Goth, C. K., Halim, A., Khetarpal, S. A., Rader, D. J., Clausen, H. & Schjoldager, K. T. 2015. A systematic study of modulation of ADAM-mediated ectodomain shedding by site-specific O-glycosylation. Proc Natl Acad Sci U S A, 112, 14623–8.

Goth, C. K., Tuhkanen, H. E., Khan, H., Lackman, J. J., Wang, S., Narimatsu, Y., Hansen, L. H., Overall, C. M., Clausen, H., Schjoldager, K. T. & Petaja-Repo, U. E. 2017. Site-specific O-Glycosylation by Polypeptide N-Acetylgalactosaminyltransferase 2 (GalNAc-transferase T2) Co-regulates beta1-Adrenergic Receptor N-terminal Cleavage. J Biol Chem, 292, 4714–4726.

Gupte, A. A., Sabek, O. M., Fraga, D., Minze, L. J., Nishimoto, S. K., Liu, J. Z., Afshar, S., Gaber, L., Lyon, C. J., Gaber, A. O. & Hsueh, W. A. 2014. Osteocalcin protects against nonalcoholic steatohepatitis in a mouse model of metabolic syndrome. Endocrinology, 155, 4697–705.

Hannemann, A., Friedrich, N., Spielhagen, C., Rettig, R., Ittermann, T., Nauck, M. & Wallaschofski, H. 2013. Reference intervals for serum osteocalcin concentrations in adult men and women from the study of health in Pomerania. BMC Endocr Disord, 13, 11.

Hansen, L. H., Madsen, T. D., Goth, C. K., Clausen, H., Chen, Y., Dzoyashvili, N., Iyer, S. R., Sangaralingham, S. J., Burnett, J. C., Jr., Rehfeld, J. F., Vakhrushev, S. Y., Schjoldager, K. T. & Goetze, J. P. 2019. Correction: Discovery of O-glycans on atrial natriuretic peptide (ANP) that affect both its proteolytic degradation and potency at its cognate receptor. J Biol Chem, 294, 18516.

Kato, K., Jeanneau, C., Tarp, M. A., Benet-Pages, A., Lorenz-Depiereux, B., Bennett, E. P., Mandel, U., Strom, T. M. & Clausen, H. 2006. Polypeptide GalNAc-transferase T3 and familial tumoral calcinosis. Secretion of fibroblast growth factor 23 requires O-glycosylation. J Biol Chem, 281, 18370–7.

Khrimian, L., Obri, A., Ramos-Brossier, M., Rousseaud, A., Moriceau, S., Nicot, A. S., Mera, P., Kosmidis, S., Karnavas, T., Saudou, F., Gao, X. B., Oury, F., Kandel, E. & Karsenty, G. 2017. Gpr158 mediates osteocalcin’s regulation of cognition. J Exp Med, 214, 2859–2873.

Kingsley, D. M., Kozarsky, K. F., Hobbie, L. & Krieger, M. 1986. Reversible defects in O-linked glycosylation and LDL receptor expression in a UDP-Gal/UDP-GalNAc 4-epimerase deficient mutant. Cell, 44, 749–59.

Kosmidis, S., Polyzos, A., Harvey, L., Youssef, M., Denny, C. A., Dranovsky, A. & Kandel, E. R. 2018. RbAp48 Protein Is a Critical Component of GPR158/OCN Signaling and Ameliorates Age-Related Memory Loss. Cell Rep, 25, 959–973 e6.

Lacombe, J., Al Rifai, O., Loter, L., Moran, T., Turcotte, A. F., Grenier-Larouche, T., Tchernof, A., Biertho, L., Carpentier, A. C., Prud’Homme, D., Rabasa-Lhoret, R., Karsenty, G., Gagnon, C., Jiang, W. & Ferron, M. 2020. Measurement of bioactive osteocalcin in humans using a novel immunoassay reveals association with glucose metabolism and beta-cell function. Am J Physiol Endocrinol Metab, 318, E381–E391.

Lacombe, J., Karsenty, G. & Ferron, M. 2013. In vivo analysis of the contribution of bone resorption to the control of glucose metabolism in mice. Mol Metab, 2, 498-504.

Lee, K. N., Jackson, K. W., Christiansen, V. J., Dolence, E. K. & Mckee, P. A. 2011. Enhancement of fibrinolysis by inhibiting enzymatic cleavage of precursor alpha2-antiplasmin. J Thromb Haemost, 9, 987–96.

Lee, K. N., Jackson, K. W., Christiansen, V. J., Lee, C. S., Chun, J. G. & Mckee, P. A. 2006. Antiplasmin-cleaving enzyme is a soluble form of fibroblast activation protein. Blood, 107, 1397–404.

Lin, X., Brennan-Speranza, T. C., Levinger, I. & Yeap, B. B. 2018. Undercarboxylated Osteocalcin: Experimental and Human Evidence for a Role in Glucose Homeostasis and Muscle Regulation of Insulin Sensitivity. Nutrients, 10.

Madsen, T. D., Hansen, L. H., Hintze, J., Ye, Z., Jebari, S., Bjoerklund Andersen, D., Joshi, H. J., Ju, T., Goetze, J. P., Martin, C., Rosenkilde, M. M., Holst, J. J., Kuhre, R. E., Goth, C. K., Vakhrushev, S. Y. & Schjoldager, K. T. 2020. An atlas of O-linked glycosylation on peptide hormones reveals diverse biological roles. Nat Commun, In press.

May, P., Bock, H. H., Nimpf, J. & Herz, J. 2003. Differential glycosylation regulates processing of lipoprotein receptors by gamma-secretase. J Biol Chem, 278, 37386–92.

Mera, P., Ferron, M. & Mosialou, I. 2018. Regulation of Energy Metabolism by Bone-Derived Hormones. Cold Spring Harb Perspect Med, 8.

Mera, P., Laue, K., Ferron, M., Confavreux, C., Wei, J., Galan-Diez, M., Lacampagne, A., Mitchell, S. J., Mattison, J. A., Chen, Y., Bacchetta, J., Szulc, P., Kitsis, R. N., De Cabo, R., Friedman, R. A., Torsitano, C., Mcgraw, T. E., Puchowicz, M., Kurland, I. & Karsenty, G. 2016a. Osteocalcin Signaling in Myofibers Is Necessary and Sufficient for Optimum Adaptation to Exercise. Cell Metab, 23, 1078–92.

Mera, P., Laue, K., Wei, J., Berger, J. M. & Karsenty, G. 2016b. Osteocalcin is necessary and sufficient to maintain muscle mass in older mice. Mol Metab, 5, 1042–7.

Morell, A. G., Gregoriadis, G., Scheinberg, I. H., Hickman, J. & ASHWELL, 1971. The role of sialic acid in determining the survival of glycoproteins in the circulation. J Biol Chem, 246, 1461–7.

Narimatsu, Y., Joshi, H. J., Nason, R., Van Coillie, J., Karlsson, R., Sun, L., Ye, Z., Chen, Y. H., Schjoldager, K. T., Steentoft, C., Furukawa, S., Bensing, B. A., Sullam, P. M., Thompson, A. J., Paulson, J. C., Bull, C., Adema, G. J., Mandel, U., Hansen, L., Bennett, E. P., Varki, A., Vakhrushev, S. Y., Yang, Z. & Clausen, H. 2019a. An Atlas of Human Glycosylation Pathways Enables Display of the Human Glycome by Gene Engineered Cells. Mol Cell, 75, 394–407 e5.

Narimatsu, Y., Joshi, H. J., Schjoldager, K. T., Hintze, J., Halim, A., Steentoft, C., Nason, R., Mandel, U., Bennett, E. P., Clausen, H. & Vakhrushev, S. Y. 2019b. Exploring Regulation of Protein O-Glycosylation in Isogenic Human HEK293 Cells by Differential O-Glycoproteomics. Mol Cell Proteomics, 18, 1396–1409.

Novak, J. F., Hayes, J. D. & Nishimoto, S. K. 1997. Plasmin-mediated proteolysis of osteocalcin. J Bone Miner Res, 12, 1035–42.

Oury, F., Ferron, M., Huizhen, W., Confavreux, C., Xu, L., Lacombe, J., Srinivas, P., Chamouni, A., Lugani, F., Lejeune, H., Kumar, T. R., Plotton, I. & Karsenty, G. 2013a. Osteocalcin regulates murine and human fertility through a pancreas-bone-testis axis. J Clin Invest, 123, 2421–33.

Oury, F., Khrimian, L., Denny, C. A., Gardin, A., Chamouni, A., Goeden, N., Huang, Y. Y., Lee, H., Srinivas, P., Gao, X. B., Suyama, S., Langer, T., Mann, J. J., Horvath, T. L., Bonnin, A. & Karsenty, G. 2013b. Maternal and offspring pools of osteocalcin influence brain development and functions. Cell, 155, 228–41.

Oury, F., Sumara, G., Sumara, O., Ferron, M., Chang, H., Smith, C. E., Hermo, L., Suarez, S., Roth, B. L., Ducy, P. & Karsenty, G. 2011. Endocrine Regulation of Male Fertility by the Skeleton. Cell, 144, 796–809.

Paczek, L., Michalska, W. & Bartlomiejczyk, I. 2008. Trypsin, elastase, plasmin and MMP-9 activity in the serum during the human ageing process. Age Ageing, 37, 318–23.

Paczek, L., Michalska, W. & Bartlomiejczyk, I. 2009. Proteolytic enzyme activity as a result of aging. Aging Clin Exp Res, 21, 9–13.

Perlman, S., Van Den Hazel, B., Christiansen, J., Gram-Nielsen, S., Jeppesen, C. B., Andersen, K. V., Halkier, T., Okkels, S. & Schambye, T. 2003. Glycosylation of an N-terminal extension prolongs the half-life and increases the in vivo activity of follicle stimulating hormone. J Clin Endocrinol Metab, 88, 3227–35.

Perrine, C. L., Ganguli, A., Wu, P., Bertozzi, C. R., Fritz, T. A., Raman, J., Tabak, L. A. & Gerken, T. A. 2009. Glycopeptide-preferring polypeptide GalNAc transferase 10 (ppGalNAc T10), involved in mucin-type O-glycosylation, has a unique GalNAc-O-Ser/Thr-binding site in its catalytic domain not found in ppGalNAc T1 or T2. J Biol Chem, 284, 20387–97.

Pi, M., Wu, Y. & Quarles, L. D. 2011. GPRC6A Mediates Responses to Osteocalcin in beta-Cells In Vitro and Pancreas In Vivo. J Bone Miner Res, 26, 1680–3.

Rawlings, N. D., Morton, F. R., Kok, C. Y., Kong, J. & Barrett, A. J. 2008. MEROPS: the peptidase database. Nucleic Acids Res, 36, D320–5.

Runkel, L., Meier, W., Pepinsky, R. B., Karpusas, M., Whitty, A., Kimball, K., Brickelmaier, M., Muldowney, C., Jones, W. & Goelz, S. E. 1998. Structural and functional differences between glycosylated and non-glycosylated forms of human interferon-beta (IFN-beta). Pharm Res, 15, 641–9.

Saavedra, Y. G., Zhang, J. & Seidah, N. G. 2013. PCSK9 prosegment chimera as novel inhibitors of LDLR degradation. PLoS One, 8, e72113.

Sabek, O. M., Nishimoto, S. K., Fraga, D., Tejpal, N., Ricordi, C. & Gaber, A. O. 2015. Osteocalcin Effect on Human beta-Cells Mass and Function. Endocrinology, 156, 3137–46.

Sanchez-Garrido, M. A., Habegger, K. M., Clemmensen, C., Holleman, C., Muller, T. D., Perez-Tilve, D., Li, P., Agrawal, A. S., Finan, B., Drucker, D. J., Tschop, M. H., Dimarchi, R. D. & Kharitonenkov, A. 2016. Fibroblast activation protein (FAP) as a novel metabolic target. Mol Metab, 5, 1015–1024.

Schjoldager, K. T. & Clausen, H. 2012. Site-specific protein O-glycosylation modulates proprotein processing - deciphering specific functions of the large polypeptide GalNAc-transferase gene family. Biochim Biophys Acta, 1820, 2079–94.

Schjoldager, K. T., Vakhrushev, S. Y., Kong, Y., Steentoft, C., Nudelman, A. S., Pedersen, N. B., Wandall, H. H., Mandel, U., Bennett, E. P., Levery, S. B. & Clausen, H. 2012. Probing isoform-specific functions of polypeptide GalNAc-transferases using zinc finger nuclease glycoengineered SimpleCells. Proc Natl Acad Sci U S A, 109, 9893–8.

Schjoldager, K. T., Vester-Christensen, M. B., Bennett, E. P., Levery, S. B., Schwientek, T., Yin, W., Blixt, O. & Clausen, H. 2010. O-glycosylation modulates proprotein convertase activation of angiopoietin-like protein 3: possible role of polypeptide GalNAc-transferase-2 in regulation of concentrations of plasma lipids. J Biol Chem, 285, 36293–303.

Steentoft, C., Vakhrushev, S. Y., Joshi, H. J., Kong, Y., Vester-Christensen, M. B., Schjoldager, K. T., Lavrsen, K., Dabelsteen, S., Pedersen, N. B., Marcos-Silva, L., Gupta, R., Bennett, E. P., Mandel, U., Brunak, S., Wandall, H. H., Levery, S. B. & Clausen, H. 2013. Precision mapping of the human O-GalNAc glycoproteome through SimpleCell technology. EMBO J, 32, 1478–88.

Steentoft, C., Vakhrushev, S. Y., Vester-Christensen, M. B., Schjoldager, K. T., Kong, Y., Bennett, E. P., Mandel, U., Wandall, H., Levery, S. B. & Clausen, H. 2011. Mining the O-glycoproteome using zinc-finger nuclease-glycoengineered SimpleCell lines. Nat Methods, 8, 977–82.

Tamura, Y., Kawao, N., Okada, K., Yano, M., Okumoto, K., Matsuo, O. & Kaji, H. 2013. Plasminogen activator inhibitor-1 is involved in streptozotocin-induced bone loss in female mice. Diabetes, 62, 3170–9.

Turcotte, A. F., Grenier-Larouche, T., Lacombe, J., Carreau, A. M., Carpentier, A. C., Mac-Way, F., Tchernof, A., Richard, D., Biertho, L., Lebel, S., Marceau, S., Ferron, M. & Gagnon, C. 2020. Association between changes in bioactive osteocalcin and glucose homeostasis after biliopancreatic diversion. Endocrine.

Zhou, B., Li, H., Xu, L., Zang, W., Wu, S. & Sun, H. 2013. Osteocalcin reverses endoplasmic reticulum stress and improves impaired insulin sensitivity secondary to diet-induced obesity through nuclear factor-kappaB signaling pathway. Endocrinology, 154, 1055–68.

Ziltener, H. J., Clark-Lewis, I., Jones, A. T. & Dy, M. 1994. Carbohydrate does not modulate the in vivo effects of injected interleukin-3. Exp Hematol, 22, 1070–5.

